# Layer 6A pyramidal cells subtypes form synaptic microcircuits with distinct functional and structural properties

**DOI:** 10.1101/2021.01.13.426506

**Authors:** Danqing Yang, Guanxiao Qi, Dirk Feldmeyer

## Abstract

Neocortical layer 6 plays a crucial role in sensorimotor coordination and integration through functionally segregated circuits linking intracortical and subcortical areas. However, because of the high neuronal heterogeneity and sparse intralaminar connectivity data on the cell-type specific synaptic microcircuits in layer 6 remain few and far between. To address this issue, whole-cell recordings combined with morphological reconstructions have been used to identify morpho-electric types of layer 6A pyramidal cells (PCs) in rat barrel cortex. Cortico-thalamic (CT), cortico-cortical (CC) and cortico-claustral (CCla) pyramidal cells have been distinguished based on to their distinct dendritic and axonal morphologies as well as their different electrophysiological properties. Here we demonstrate that these three types of layer 6A pyramidal cells innervate neighboring excitatory neurons with distinct synaptic properties: CT PCs establish weak facilitating synapses to other L6A PCs; CC PCs form synapses of moderate efficacy; while synapses made by putative CCla PCs display the highest release probability and a marked short-term depression. Furthermore, for excitatory-inhibitory synaptic connections in layer 6 we were able to show that both the presynaptic PC type and the postsynaptic interneuron type govern the dynamic properties of the of the respective synaptic connections. We have identified a functional division of local layer 6A excitatory microcircuits which may be responsible of the differential temporal engagement of layer 6 feed-forward and feedback networks. Our results provides a basis for further investigations on the long-range cortico-cortical, cortico-thalamic and cortico-claustral pathways.

## Introduction

Early born excitatory neurons in the ventricular zone migrate into the cortical plate to occupy deep layers; layer 6 (L6) is the first neocortical layer to form (Rakic 2009; Lodato and Arlotta 2015). In rat barrel cortex, layer 6 is the thickest layer and contains the highest number of neurons (Hutsler et al. 2005; Meyer et al. 2010). Pyramidal cells (PCs) in this layer 6 of the neocortex display a high degree of morphological, electrophysiological and molecular diversity (Zhang and Deschenes 1997; Kumar and Ohana 2008; Thomson 2010; Marx and Feldmeyer 2013; Gouwens et al. 2019; Kast et al. 2019; Egger et al. 2020; Gouwens et al. 2020). They project either *intra-telencephalically* (IT) within the cortex, to the striatum, and the claustrum or *extra-telencephalically* (ET) to e.g. different thalamic nuclei (for a review see Rockland 2019). This heterogeneity makes it difficult to elucidate the exact functional and structural properties of the L6A synaptic microcircuitry. Compared to PCs in other neocortical layers, L6 PCs rarely establish intralaminar synaptic contacts and if so, they generally display a low synaptic release probability (Beierlein and Connors 2002; Mercer et al. 2005; West et al. 2006; Crandall et al. 2017; Seeman et al. 2018). Hence, the excitatory synaptic microcircuits of layer 6 and their dynamic properties remain so far not well understood.

As the pre-eminent source of cortico-thalamic (CT) projections, L6 microcircuits provide contextual modulation in the feed-back loop of sensory processing system (Harris and Mrsic-Flogel 2013; Velez-Fort et al. 2014). The two major types of L6 principal cells, the ET CT PCs and the IT cortico-cortical (CC) show distinct axonal projection patterns and participate in distinct microcircuits within the neocortical network (Kumar and Ohana 2008; Pichon et al. 2012; Sundberg et al. 2018). CT PCs are known to generate weak and facilitatory excitatory postsynaptic responses onto both excitatory and inhibitory neurons (West et al. 2006; Frandolig et al. 2019). Conversely, L6A CC PCs have been proposed to innervate L6A PCs and parvalbumin (PV) positive interneurons; these synapses exhibit short-term synaptic depression (Mercer et al. 2005; Yang et al. 2020). Apart from L6A CC PCs another class of IT L6A PCs exists that shows axonal projections predominantly to the ipsilateral claustrum (CCla PCs). The claustrum itself is reciprocally connected with most neocortical areas and targets all cortical laminae although connections with sensory cortices are generally weaker than those with more frontal cortical regions (Zakiewicz et al. 2014; Atlan et al. 2017; Zingg et al. 2018; Rockland 2019; Gouwens et al. 2020). Among other functions such as the regulation of attention the claustrum is considered to coordinate sensory and motor modalities from different cortical areas (Naghavi et al. 2007; Smith et al. 2017; Bayat et al. 2018; Zingg et al. 2018; for a review see Jackson et al. 2020). In humans, the claustrum together with the striatum (also a part of the telencephalon) participates in a salience network which is known to integrate sensory, emotional, and cognitive information (Peters et al. 2016; Smith et al. 2019b).

Therefore, it is important to study the role of IT PCs in driving specific intra-cortical networks and conveying subcortical output. To systematically investigate the L6A excitatory microcircuitry we identified these three types of L6A PCs in rat barrel cortex by performing patch-clamp recordings and morphological reconstructions. We found that CT, CC and putative CCla PCs show distinct morphological and electrophysiological properties, allowing a clear classification of the different L6A PC types. Using paired-recordings we found that L6A excitatory neurons are differentially innervated by intralaminar PCs with respect to the presynaptic PC type. More specifically, L6A CT PCs innervate other L6A PCs; the synapses established between them have a low release probability and exhibit short-term facilitation. Synapses made by presynaptic CC PC are generally stronger and those established by putative CCla PCs display the highest release probability and a marked short-term depression. Moreover, we demonstrate that in excitatory-inhibitory synaptic connections both presynaptic PC type and postsynaptic interneuron type govern the synaptic release properties. Our results reveal a neuronal cell type-specific and functionally distinct organization of L6A excitatory microcircuits in rat barrel cortex.

## Materials and Methods

### Slice Preparation

All experimental procedures involving animals were performed in accordance with the guidelines of the Federation of European Laboratory Animal Science Association (FELASA), the EU Directive 2010/63/EU, and the German animal welfare law. In this study, Wistar rats (Charles River, either sex) aged 17–21 postnatal days were anaesthetized with isoflurane at a concentration < 0.1% and decapitated. The brain was quickly removed and placed in an ice-cold artificial cerebrospinal fluid (ACSF) containing a high Mg^2+^ and a low Ca^2+^ concentration (4 mM MgCl_2_ and 1 mM CaCl_2_) to reduce potentially excitatotoxic synaptic transmission during slicing. In order to maintain adequate oxygenation and a physiological pH level, the solution was constantly bubbled with carbogen gas (95% O_2_ and 5% CO_2_). Thalamocortical slices were cut at 350 μm thickness using a high vibration frequency and were then transferred to an incubation chamber for a recovery period of 30-60 minutes at room temperature.

During whole-cell patch-clamp recordings, slices were continuously superfused (perfusion speed ~5 ml/min) with ACSF containing (in mM): 125 NaCl, 2.5 KCl, 1.25 NaH_2_PO_4_, 1 MgCl_2_, 2 CaCl_2_, 25 NaHCO_3_, 25 D-glucose, 3 mho-inositol, 2 sodium pyruvate and 0.4 ascorbic acid, bubbled with carbogen gas and maintained at 30-33 °C. Patch pipettes were pulled from thick-wall borosilicate glass capillaries and filled with an internal solution containing (in mM): 135 K-gluconate, 4 KCl, 10 HEPES, 10 phosphocreatine, 4 Mg-ATP, and 0.3 GTP (pH 7.4 with KOH, 290-300 mOsm). The ‘searching’ pipette (see below) was filled with an internal solution in which K^+^ is replaced by Na+ (containing (in mM): 105 Na-gluconate, 30 NaCl, 10 HEPES, 10 phosphocreatine, 4 Mg-ATP and 0.3 GTP), in order to prevent the depolarization of neurons during searching for a presynaptic cell. Biocytin was added to the internal solution at a concentration of 3-5 mg/ml in order to stain patched neurons; A recording time longer than 15 minutes was necessary for an adequate diffusion of biocytin into dendrites and axons of targeted cells (Marx et al. 2012).

### Electrophysiological recordings

Neurons were visualized using infrared differential interference contrast microscopy. The barrels can be identified in layer 4 as dark stripes with light ‘hollows’ and were visible in 6-8 consecutive slices (Agmon and Connors 1991; Feldmeyer et al. 1999). L6A neurons were recorded in the upper 65% of layer 6 while neurons in L6B (Woo et al. 1991; Clancy and Cauller 1999; lower 35%; Marx and Feldmeyer 2013) were not used for recordings. Putative PCs and interneurons were differentiated on the basis of their intrinsic action potential firing pattern during recording and after post-hoc histological staining also by their morphological appearance.

Whole-cell patch clamp recordings were made using an EPC10 amplifier (HEKA, Lambrecht, Germany). Signals were sampled at 10 kHz, filtered at 2.9 kHz using Patchmaster software (HEKA), and later analyzed off-line using Igor Pro software (Wavemetrics, USA). The recordings were performed using patch pipettes of resistance between 6 and 10 MΩ. Because the intralaminar connectivity ratio in L6A is low, we performed a ‘searching procedure’ described previously after patching a postsynaptic neuron (Feldmeyer et al. 1999; Feldmeyer and Radnikow 2016). No biocytin was added to the Na-based internal solution of ‘searching’ pipettes used to identify potential synaptic connections. When an action potentials (APs) elicited in ‘loose cell-attached’ mode resulted in an excitatory postsynaptic potential (EPSP) in the postsynaptic neuron, this presynaptic neuron was re-patched with a new pipette filled with biocytin-containing internal solution.

### Histological staining

After single cell or paired-recordings, brain slices containing biocytin-filled neurons were fixed for at least 24 h at 4 °C in 100 mM phosphate buffer solution (PBS, pH 7.4) containing 4% paraformaldehyde (PFA). After rinsing slices several times in 100 mM PBS, they were treated with 1% H_2_O_2_ in PBS for about 20 min in order to reduce any endogenous peroxidase activity. Slices were rinsed repeatedly with PBS and then incubated in 1% avidin-biotinylated horseradish peroxidase (Vector ABC staining kit, Vector Lab. Inc., Burlingame, USA) containing 0.1% Triton X-100 for 1 h at room temperature. The reaction was catalysed using 0.5 mg/ml 3,3-diaminobenzidine (DAB; Sigma-Aldrich, St.Louis, Mo, USA) as a chromogen. Slices were then rinsed with 100 mM PBS, followed by slow dehydration with ethanol in increasing concentrations and finally in xylene for 2–4 h. After that, slices were embedded using Eukitt medium (Otto Kindler GmbH, Freiburg, Germany).

In a subset of experiments, we tried to identify the expression of molecular marker forkhead box protein P2 (FoxP2) in L6A PCs recorded in acute brain slices to investigate a possible correlation with the electrophysiological and morphological properties. To this end, during electrophysiological recordings, Alexa Fluor 594 dye (1:500, Invitrogen) was added to the internal solution for *post hoc* identification of patched neurons. After recording, slices (350 μm) were fixed with 4% PFA in 100 mM phosphate buffered saline (PBS) for at least 24 h at 4°C and then permeabilized in 1% milk power solution containing 0.5% Triton X-100 and 100 mM PBS. Primary and secondary antibodies were diluted in the permeabilization solution (0.5% Triton X-100 and 100 mM PBS) shortly before experiments. For single-cell FoxP2 staining, slices were incubated overnight with Goat-anti-FoxP2 primary antibody (1:500, Santa Cruz Biotechnology) at 4°C and then rinsed thoroughly with 100 mM PBS. Subsequently, slices were treated with Alexa Fluor secondary antibodies (1:500) for 2–3 h at room temperature in the dark. After being rinsed in 100 mM PBS, the slices were embedded in Fluoromount. The fluorescence images were taken using the Olympus CellSens platform. The position of the patched neurons was identified by the conjugated Alexa dye, so that the expression of FoxP2 could be tested in biocytin-stained neurons. After acquiring fluorescent images, slices were incubated in 100 mM PBS overnight and were processed for subsequent histological processing as described above.

### Morphological 3D Reconstructions

Computer-assisted morphological 3D reconstructions of biocytin-filled L6A neurons were made using NEUROLUCIDA^®^ software (MicroBrightField, Williston, VT, USA) and Olympus BV61 microscopy at 1000x magnification (100x objective, 10x eyepiece). Neurons were selected for reconstruction based on the quality of biocytin labelling when background staining was minimal. The cell body, dendritic and axonal branches were reconstructed manually under constant visual inspection to detect thin and small collaterals. Barrel and layer borders, pial surface and white matter were delineated during reconstructions at lower magnification. The position of soma and layers were confirmed by superimposing the differential interference contrast images taken during the recording. The tissue shrinkage was corrected using correction factors of 1.1 in the *x*–*y* direction and 2.1 in the *z* direction (Marx et al. 2012). Analysis of 3D reconstructed neurons was done with NEUROEXPLORER^®^ software (MicroBrightField Inc., Willston, VT, USA). Putative synaptic contacts were identified as close appositions of presynaptic axon terminals and postsynaptic dendrites in the same focal plane under light microscopy with 100× objective and 10× eyepiece. The distance between the soma and a putative synaptic contact was calculated as the path length along the dendrite from the location of synaptic contact to soma in 3D space.

### Data analysis

Custom-written macros for Igor Pro 6 (WaveMetrics, Lake Oswego, USA) were used to analyse the recorded electrophysiological signals. Passive and active action potential properties were assessed by eliciting a series of initially hyperpolarising, followed by depolarising 1 s current pulses under current clamp configuration. Neurons with a series resistance exceeding 45 MΩ (50 MΩ for neurons from paired-recordings) or with a depolarized membrane potential (> −50 mV) after rupturing the cell membrane were excluded from data analysis.

Synaptic properties were evaluated as described previously (Feldmeyer et al. 1999; Feldmeyer et al. 2002). All uEPSP recordings were aligned to their corresponding presynaptic AP peaks and an average sweep was generated as the mean uEPSP. The EPSP amplitude was calculated as the difference between the mean baseline and maximum voltage of the postsynaptic event. The paired pulse ratio was defined as the 2^nd^/3^rd^ uEPSP divided by the 1^st^ uEPSP amplitude elicited by presynaptic APs at a stimulation frequency of 10 Hz. Failures were defined as events with an amplitude <1.5× the standard deviation (SD) of the noise within the baseline window; the failure rate refers to the percentage of failures. The coefficient of variation (CV) was calculated as the SD divided by the mean uEPSP amplitude. Rise time was calculated as the mean time to rise from 20% to 80% of the peak amplitude. Latency was calculated as the time interval between the peak amplitude and the onset of the EPSP. Decay time was measured using single exponential fit to the decay phase of both individual and averaged responses.

For all data, the mean ± SD was given. Statistical comparisons among multiple groups were done using a Kruskal-Wallis test followed by a Dunn-Holland-Wolfe non-parametric multiple comparison test. Wilcoxon Mann-Whitney U test was performed to access significant difference between individual clusters. Statistical significance was set at P < 0.05, n indicates the number of neurons/pairs analysed.

## Results

### Three types of morphologically and electrophysiologically distinct L6A pyramidal cells in rat barrel cortex

Whole-cell recordings from L6A excitatory neurons (n = 125) with simultaneous biocytin fillings were performed in acute brain slices of rat barrel cortex allowing post-hoc identification of their morphology. Neurons with incomplete filling, high background staining or major truncations of the dendrites were excluded from morphological analysis, resulting in 41 high quality 3D reconstructions of L6A PCs (**Fig. S1**). In accordance with previous studies, three major classes of L6A PCs can be distinguished by their dendritic and axonal projecting patterns (Katz 1987; Kumar and Ohana 2008; Baker et al. 2018; Cotel et al. 2018) (**Fig. 1 A, B**). CT-like PCs have an apical dendrite terminating predominately in layer 4 and axon collaterals that project daily vertically. Their basal dendrites and axons are comparatively short (1899 ± 447 μm for length of basal dendrites and 5502 ± 2189 μm for axonal length, respectively) and have a small horizontal fieldspan (231 ± 58 μm for dendritic and 529 ± 240 μm for axonal horizontal fieldspan, respectively). Dendrites of upright CC PCs resemble those of CT PCs. Their apical dendrite projects towards the pia and terminates in layer 4 or layer 5A. Basal dendrites, however, are longer than those of CT PCs and have a larger horizontal fieldspan (length: 3770 ± 1277 μm; horizontal dendritic fieldspan: 386 ± 57 μm). Consistent with previous studies, we found that CC PCs have long, horizontal projections of axons across several barrel columns (16356 ± 4081 μm for axonal length and 1545 ± 541 μm for axonal horizontal fieldspan). In addition, we identified another subpopulation of PCs exhibiting long, sparsely tufted apical dendrites that reach layer 1 and exhibit horizontally expanding basal dendrites. A similar dendritic morphology has been found for IT claustrum-projecting neurons in layer 6 of rat and cat primary visual cortex (Katz 1987; Cotel et al. 2018) and are therefore named CCla PCs in the remainder of this manuscript. Here, these putative CCla PCs were found to have the largest vertical and horizontal dendritic fieldspan of the three L6A PC subtypes **(Fig. 1C and Tab. S1)**. Moreover, it worth noting that CCla PCs have fewer basal dendrites compared to CT (3.6 ± 0.9 vs. 6.6 ± 1.6, P = 4.5E-06) and CC PCs (3.6 ± 0.9 vs. 6.3 ± 1.8, P = 4.5E-06), but each single basal dendrite exhibits more collaterals that ramify profusely (average length of basal dendrite: 917 ± 358 μm vs. 295 ± 59 μm for CT PCs and 607 ± 170 for CC PCs, p < 0.001). The axonal length (10224 ± 4648 μm) and horizontal fieldspan (1101 ± 479 μm) of CCla PCs range between those of CT and CC PCs **(Fig. 1C)**.

**Fig. 1.**
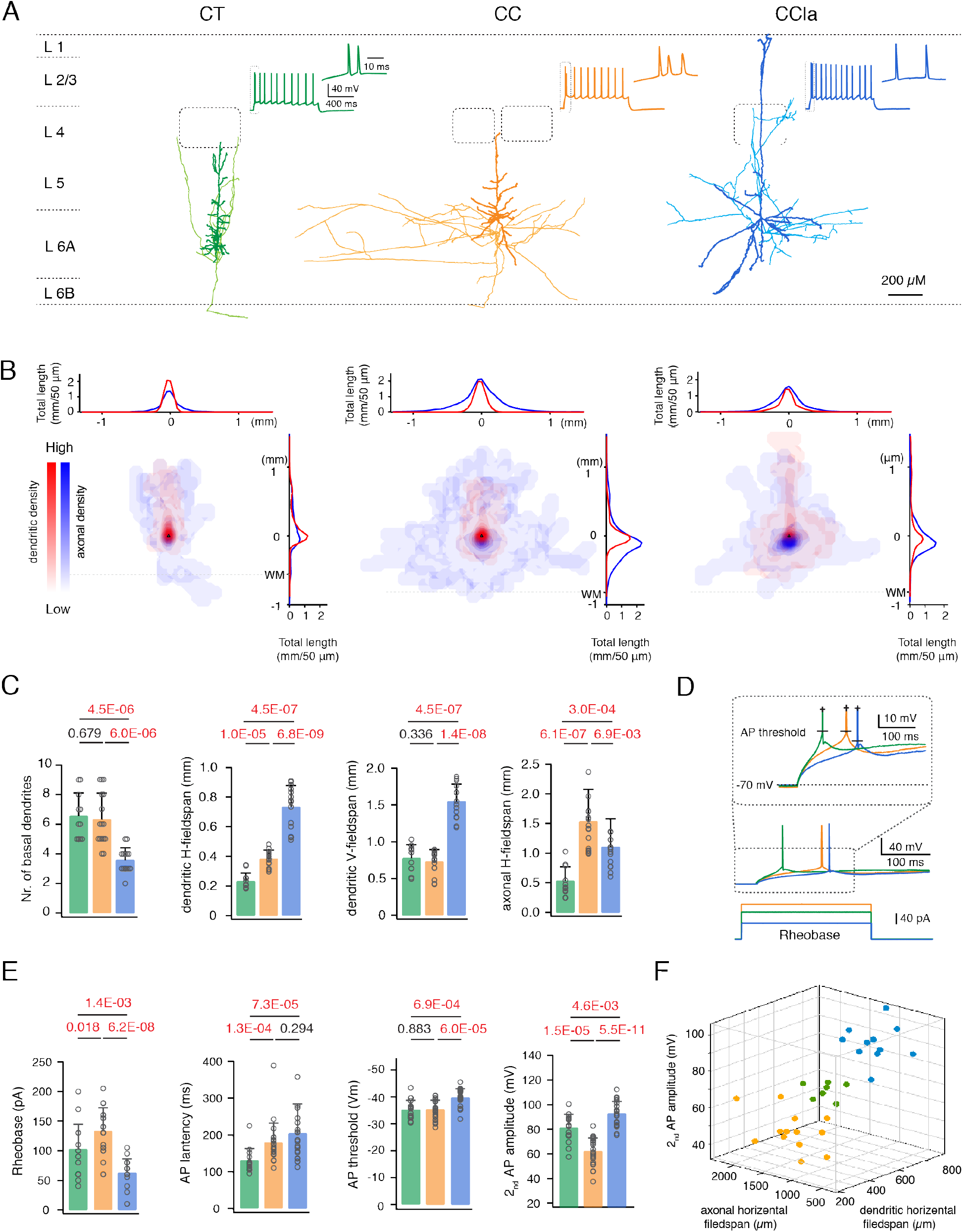
Identification of three subtypes of pyramidal cells in L6A of rat barrel cortex. **(A)** Representative morphological reconstructions of a CT (green) PC, a CC PC (orange) and a putative CCla PC (blue); the somatodendritic domain is given in a darker, the axons in a lighter shade). Inset, the corresponding firing patterns in low and higher magnification. **(B)** 2D density maps of L6A CT PCs (left, n = 11), CC PCs (center, n = 16) and CCla PCs (right, n = 14). Dendrites are shown in red and axons are shown in blue. Horizontal distribution of L6 PC dendrites and axons are shown on the top while vertical distribution are shown at the right. The curves indicate the average dendritic and axonal density distribution; bin size in the x and y axis: 50 μm in horizontal and vertical direction. Dashed lines indicate white matter position. **(C)** Summary data of several morphological properties of L6A CT (n = 11), CC (n = 16) and CCla (n = 14) PCs. Data were compared between groups and p values calculated using the Wilcoxon Mann-Whitney U test. **(D)** 3D scatter plot showing a clear separation of three L6A PC subtypes using morphological and electrophysiological properties. CT PCs in green, CC PCs in orange and CCla PCs in blue. **(E)** Summary data of several electrophysiological properties of L6A CT (n = 15), CC (n = 23) and CCla (n = 22) PCs. To calculate p values the Wilcoxon Mann-Whitney U test was used. **(F)** First elicited AP (middle) of a representative CT (green), CC (orange) and CCla (blue) PC in response to rheobase current injection (bottom). Higher magnification of first APs (inset) illustrating the difference in AP threshold and latency.

In brain slice preparations, long-range axonal collaterals of CC and CCla PCs are massively truncated and consequently the length and fieldspan estimates are strongly underestimated (s. e.g. Egger et al. 2020). Despite this, the distinct axonal and dendritic properties of the three L6A PC types allow an unambiguous cell type identification. Density plots illustrating dendritic and axonal distribution of each PC type are shown in **Fig. 1 B** and individual reconstruction are shown in **Fig. S1**. Additionally, we measured the relative distance form soma to pial surface to analyse relationship between the soma laminar location and morphological patterns. We found that L6A CC and CCla PCs were located closer to the border between layers 5B and 6A while CT PCs distribute preferentially in deeper L6A. More details regarding morphological properties and statistical comparison are shown in **Tab. S1**.

In addition to their different morphological properties, L6A CT and CC PCs have been found to exhibit distinct passive and active electrophysiological characteristics (Kumar and Ohana 2008; Tian et al. 2014). Here, we systematically analysed and compared the intrinsic electrophysiological properties of morphologically identified CT, CC and putative CCla PCs. When compared to CC PCs, CT PCs showed a significantly higher input resistance and a longer onset time for the first AP evoked by a lower rheobase current. CC PCs generally exhibit an initial spike burst composed of doublet or triplet riding on a depolarizing envelope and display a larger first and second AP half-width **(Fig. 1 A, Tab. S1)**. CCla PCs showed a similar regular firing pattern without an initial spike burst as found for CT PCs, but with a significantly larger AP amplitude than that of the other two PC types (102 ± 7 mV for CCla PCs, 93 ± 8 mV for CT PCs and 92 ± 7 mV for CC PCs). Furthermore, putative CCla PCs showed the highest input resistance (281 ± 73 MΩ for CCla PCs, 235 ± 80 MΩ for CT PCs and 152 ± 35 MΩ for CC PCs) and the most hyperpolarized AP threshold potential (−39.8 ± 3.3 mV for CCla PCs, −35.1 ± 3.5 for CT PCs and −35.2 ± 3.5 for CC PCs) by injecting the lowest rheobase current (63 ± 23 pA for CCla PCs, 102 ± 43 for CT PCs and 133 ± 39 for CC PCs) **(Fig.1 D, E)**. This indicates that CCla PCs have a higher membrane excitability than the other L6A PC subclasses. Therefore, our results demonstrate that L6A PCs can be reliably discriminated and classified on the basis of both morphological and physiological features as shown by the 3D scatter plot in **Fig. 1F**. More electrophysiological properties and the statistical comparison of the three L6A PC types are shown in **Tab. S1**.

The nuclear transcription factor FoxP2 has been shown to be a molecular marker for CT L6A PCs (Hisaoka et al. 2010; Sundberg et al. 2018). To identify the expression of FoxP2 in L6A PCs, we performed whole-cell recordings with simultaneous filling of biocytin and the fluorescent Alexa Fluor 594 dye. Subsequently, brain slices were processed for FoxP2 immunofluorescence staining. We found that the morphological and electrophysiological identified CT L6A PCs are FoxP2-positive, while both CC PCs and putative CCla PCs are FoxP2-negative **(Fig. S2)**. The correlation between neuronal morphology, electrophysiology, and FoxP2 expression demonstrates the reliability of our cell type classification.

### Specific synaptic properties of presynaptic CT, CC and CCla PCs innervating other L6A PCs

Because of the low excitatory synaptic connectivity ratio in layer 6A, a so called ‘loose seal’ searching protocol (see ‘Materials and Methods’) was used to test for potential synaptic connections. 194 excitatory neurons were recorded as postsynaptic neurons and 1513 potential presynaptic neurons were tested. Out of the tested cells 96 were found to be synaptically coupled with the recorded excitatory neurons, so that the connectivity ratio of L6A excitatory to excitatory (E→E) cell pairs was 6.3%. It should be noted however, that in acute brain slices axonal collaterals are have a high degree of truncation resulting in an underestimate of the actual synaptic connectivity.

Overall, 47 cell pairs were recorded successfully in dual whole-cell mode. After post-hoc morphological reconstructions, the pre- and postsynaptic PC type were identified according to their specific features shown in **Fig. 1**. As in previous studies, the L6A bipolar, multipolar and inverted excitatory neurons were classified as CC PCs based on their horizontal axonal morphology and initial burst-spiking firing behavior (Zhang and Deschenes 1997; Kumar and Ohana 2008; for a review see: Thomson 2010). There are four synaptically coupled pairs with a presynaptic CT PC (3 CT→CT, 1 CT→CC), 36 pairs with a presynaptic CC PC (23 CC→CC, 13 CC→CT) and 7 pairs with a presynaptic CCla PC (6 CCla→CC, 1 CCla→CT). Morphological reconstructions and synaptic recordings of individual synaptically coupled pairs are shown in **Fig. S3**.

As expected from their axonal projection pattern, we found that CT PCs rarely form synaptic connections with other L6A PCs as presynaptic cells (4 out of 47 pairs, 8.5%). Connections with CT PCs showed small amplitude unitary EPSPs (0.09 ± 0.10 mV) with a high CV (1.03 ± 0.25) and a high failure rate (64.4 ± 31.1 %). Eliciting three APs in a presynaptic CT PC at an inter-spike interval of 100 ms resulted invariably in a strong short-term facilitation of the unitary synaptic response, as characterized by a mean PPR2nd/1st of 2.22 ± 1.24 and PPR3rd/1st of 1.89 ± 0.91 (**Fig. 2 A, B**). In contrast to CT PCs, the vast majority of E→E connections were established by presynaptic CC PCs (36 out of 47 PCs, 76.6%). CC PCs were found to preferably innervate other CC PCs (n = 23) rather than CT PCs (n = 13), probably due to the smaller dendritic length and horizontal fieldspan of postsynaptic CT PCs (**Fig. 1 D**). The E→E connections with a presynaptic CT PC displayed uEPSPs with an average amplitude of 0.37 ± 0.24 mV, which is significantly larger than those evoked by presynaptic CT PCs. The uEPSP amplitudes at this L6A connection type varied widely (from 0.03 to 0.98 mV) and exhibited either short-term depression or facilitation, which on average resulted in a paired pulse ratio close to 1 (PPR2nd/1st: 1.19 ± 0.56, PPR3rd/1st: 1.08 ± 0.39)(**Fig. 2 A-C**); the mean CV and failure rate were 0.68 ± 0.23 and 35.5 ± 23.8 %, respectively, suggesting that CC PCs form synapses of moderate efficacy (**Fig. 2 D**). We also compared the functional properties of synaptic connections established by L6A CCla PCs. In contrast to the other two L6A PC subclasses, CCla PCs establish strong, reliable synaptic connections that displayed the largest average uEPSP amplitude (0.70 ± 0.40 mV), a comparatively small CV (0.57 ± 0.22) and a low failure rate (18.8 ± 18.9 %). In response to a 10 Hz train of three APs elicited in a presynaptic CCla PC, EPSPs displayed short-term depression with a mean PPR2nd/1st of 0.94 ± 0.25 and PPR3rd/1st of 0.74 ± 0.21 (**Fig.2 C, D**). The difference in short-term synaptic plasticity between CT, CC and CCla synaptic connections in layer 6A is even more evident in response to a train of ten presynaptic APs (**Fig. S4**). On the other hand, the EPSP time course was similar for the different L6A connection types (**Tab. 1**).

**Fig. 2.**
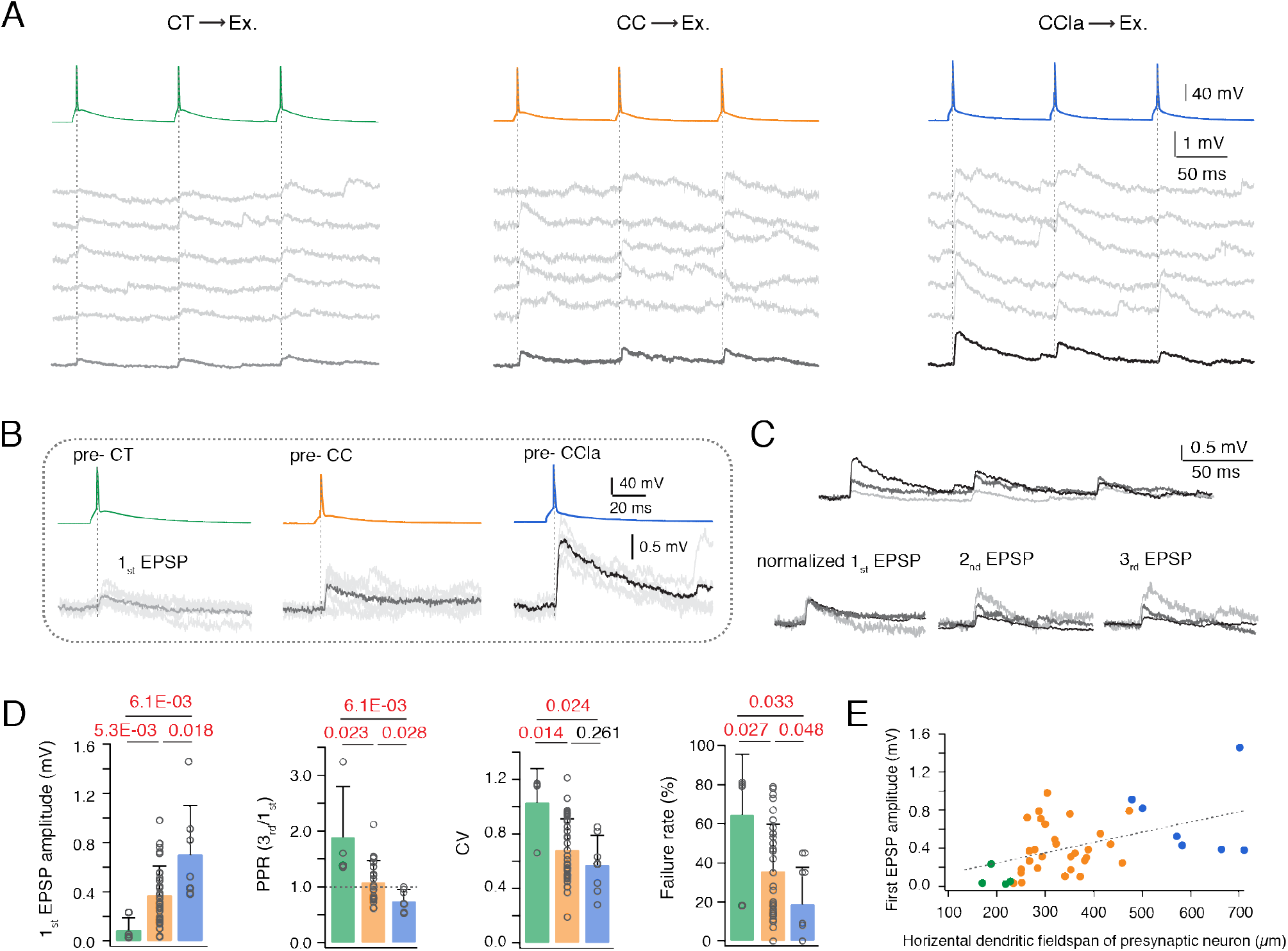
The properties of L6A E→E connections dependent of the different presynaptic PC subtypes. **(A)** Unitary synaptic connections in L6A with a presynaptic CT (left), CC (middle) and CCla (right) PC. Five consecutive EPSPs (middle) and average EPSP (bottom) are elicited by presynaptic APs (top, 10Hz). **(B)** First EPSP induced by AP of a presynaptic CT, CC and CCla PC. The mean EPSP waveform is shown in light gray for pre-CT pair, in gray for pre-CC pair and in dark gray for pre-CCla pair. **(C)** Top, overlay of average EPSPs recorded in a monosynaptic connection with a presynaptic CT (light gray), CC (gray) and CCla PC (dark gray). Bottom, normalising the mean EPSP amplitudes reveals the difference in PPR for these three connection types. **(D)** Summary data of several synaptic properties of L6A E→E connections with presynaptic CT (green, n = 4), CC (orange, n = 36) and CCla (blue, n = 7) PCs. P values are calculated using Wilcoxon Mann-Whitney U test. **(E)** Plot of 1st EPSP amplitude versus horizontal dendritic fields-pan of the presynaptic neuron in L6A E→E connections. Best linear fit is shown in gray dashed line (r = 0.47, p = 0.0014). Color coding as in (D).

**Tab. 1.** Functional and morphological properties of L6A E→E synaptic connections. Italic bold font indicates significant differences; *P < 0.05, **P < 0.01 for Wilcoxon Mann-Whitney *U* test.

|  | CT- pairs<br>(n = 4) | CC-pairs<br>(n = 36) | CCLa-pairs<br>(n = 7) | CT- vs. CC-<br>pairs | CT- vs. CCLa-<br>pairs | CC- vs. CCLa-<br>pairs |
| --- | --- | --- | --- | --- | --- | --- |
| Electrophysiological properties |  |  |  |  |  |  |
| Amplitude (mV) | 0.09 ± 0.10<br>(0.03 to 0.24) | 0.37 ± 0.24<br>(0.03 to 0.98) | 0.70 ± 0.40<br>(0.38 to 1.46) | ***\*\*0.0053*** | ***\*\*0.0061*** | ***\*0.0183*** |
| PPR (2nd/1st) | 2.22 ± 1.24<br>(1.15 to 3.73) | 1.19 ± 0.56<br>(0.62 to 3.21) | 0.94 ± 0.25<br>(0.60 to 1.36) | ***\*0.0358*** | ***\*0.0242*** | 0.0860 |
| PPR (3rd/1st) | 1.89 ± 0.91<br>(1.35 to 3.24) | 1.08 ± 0.39<br>(0.62 to 2.12) | 0.74 ± 0.21<br>(0.53 to 1.00) | ***\*0.0234*** | ***\*\*0.0061*** | ***\*0.0282*** |
| CV | 1.03 ± 0.25<br>(0.66 to 1.17) | 0.68 ± 0.23<br>(0.37 to 1.21) | 0.57 ± 0.22<br>(0.28 to 0.85) | ***\*0.0143*** | ***\*0.0242*** | 0.2609 |
| Failure rate (%) | 64.4 ± 31.1<br>(18 to 81) | 35.5 ± 23.8<br>(0 to 79) | 18.8 ± 18.9<br>(0 to 45) | ***\*0.0271*** | ***\*0.0333*** | ***\*0.0475*** |
| Rise time (ms) | 0.91 ± 0.52<br>(0.24 to 1.52) | 1.44 ± 0.99<br>(0.50 to 5.72) | 1.44 ± 0.34<br>(1.14 to 1.96) | 0.3674 | 0.0697 | 0.4438 |
| Latency (ms) | 2.37 ± 0.86<br>(0.66 to 1.17) | 1.73 ± 0.92<br>(0.80 to 4.67) | 1.75 ± 0.70<br>(0.28 to 0.85) | 0.1150 | 0.2303 | 0.5681 |
| Decay time (ms) | 34.4 ± 26.1<br>(29.0 to 54.8) | 38.8 ± 17.9<br>(12.7 to 73.1) | 37.1 ± 9.0<br>(9.4 to 61.4) | 0.6878 | 0.8750 | 0.8586 |
| Morphological properties |  |  |  |  |  |  |
| Nr. of contacts<br>per connection | 3.3 ± 2.1<br>(1 to 5) | 4.1 ± 1.8<br>(2 to 7) | 3.8 ± 1.2<br>(2 to 5) | 0.7097 | 0.8810 | 0.4706 |
| Geometric<br>distance (µm) | 65.3 ± 36.8<br>(36 to 150) | 119.0 ± 60.4<br>(30 to 239) | 149.1 ± 57.0<br>(36 to 237) | ***\*\*0.0037*** | ***\*\*\*0.0002*** | ***\*0.0377*** |
| Inter-soma<br>distance (µm) | 75.6 ± 92.3<br>(21 to 214) | 78.1 ± 67.4<br>(20 to 345) | 86.4 ± 77.1<br>(23 to 235) | 0.3004 | 0.2303 | 0.5589 |

To investigate whether synaptic properties depend also on the postsynaptic L6A PC subtype, we compared functional properties of morphologically identified CC→CC (n = 23) and CC→CT (n = 11) connections and found no significant difference in EPSP properties (**Fig. S5**). Taken together, our results suggest that functional properties of E→E connections in layer 6A are presynaptic cell-type specific but not depending on postsynaptic target. A summary plot of first EPSP amplitude versus presynaptic dendritic horizontal fieldspan is given in **Fig. 2E**, demonstrating a tight correlation between presynaptic neuron morphology and postsynaptic uEPSP properties.

We also studied the morphological characteristics of L6A synaptic connections between excitatory neurons (E→E connections). The average distance between the cell bodies of pre- and postsynaptic neurons is similar for the different connection types (**Tab. 1**) and no correlation between uEPSP amplitude and inter-soma distance was found. To assess the number of putative synaptic contacts, we searched for close appositions of presynaptic axon terminals and postsynaptic dendrites under light microscopy (**Fig. S6 A-C**). Comparison of the number of putative synaptic contacts between CT-, CC- and CCla- formed connections revealed no marked difference (**Tab. 1, Fig. S6 E**). This suggests that difference in the number of synaptic contacts are probably not responsible for the cell type-specific functional properties of L6A E→E connections. Considering the distinct short-term synaptic plasticity and difference in CV and failure rate, we conclude that presynaptic CT, CC and CCla PCs show weak, moderate and comparatively strong synaptic release probability, respectively, in synaptic connections with other L6A PCs. Moreover, light microscopic examination suggests that CT PCs established putative synaptic contacts on the proximal portion of the basal dendrites of L6A PCs with an average geometric distance of 65.3 ± 36.8 μm. For connections formed by CC PCs, putative synaptic contacts were found both on proximal and the distal portion of the basal dendrites resulting in a significantly larger synapse-to-soma distance (119.0 ± 60.4 μm) when compared with connections with a presynaptic CC PCs. L6A CCla PCs formed putative synapses on distal collaterals of basal dendrites and proximal apical oblique dendrites, displaying the largest average synapse-to-soma distance of 149.1 ± 57.0 μm among the three L6A E→E connection types (**Fig. S6 D, E**). These data indicate that although L6A E→E connections show similar EPSP time course (**Tab. 1**), the precise location of synapses on postsynaptic dendrites is specific to the presynaptic cell type.

### Characterization of pyramidal cell to interneuron connections in layer 6A of rat barrel cortex

Cortical GABAergic interneurons show a highly diverse firing pattern which depends largely on the interneuron type (Gupta et al. 2000; Ascoli et al. 2008; Yuste et al. 2020). Fast spiking (FS) interneurons generate high frequency APs without apparent frequency accommodation. The remaining interneurons are the so-called non fast-spiking (nFS) neurons which comprise a large group of irregular-spiking, late spiking, and burst spiking interneurons (Kawaguchi and Kubota 1996; Defelipe et al. 2013; Emmenegger et al. 2018). Both FS and nFS interneurons are broad families with different transcriptomic, electrophysiological and morphological phenotypes (Gouwens et al. 2020; Scala et al. 2020; Yuste et al. 2020). Excitatory synapses onto FS interneurons are initially strong (i.e. have a high synaptic release probability) and depress with ongoing stimulation; excitatory synapses onto nFS neurons are generally weak (i.e. have a low synaptic release probability) and show facilitation upon repetitive firing (Tan et al. 2008; Caputi et al. 2009). In order to comprehensively investigate synaptic microcircuits between a L6A excitatory neuron and an inhibitory interneuron (E→I connection), we broadly classified L6A interneurons into a FS (n = 23) and a nFS (n = 30) group of interneurons on the basis of their electrophysiological properties. The characteristic feature of L6A FS interneurons was a high frequency firing behavior with a low frequency adaptation; nFS interneurons, on the other hand, were characterized by a low rheobase, high adaptation and a lower firing frequency (**Fig. 3A, B**). Moreover, L6A FS and nFS interneurons also display significant differences in AP half-width (HW) and afterhyperpolarization (AHP) amplitude (**Fig. 3C**). The 3D scatter plots in **Fig. 3D** illustrate the reliability of the electrophysiological classification of these two L6A interneuron groups.

**Fig. 3.**
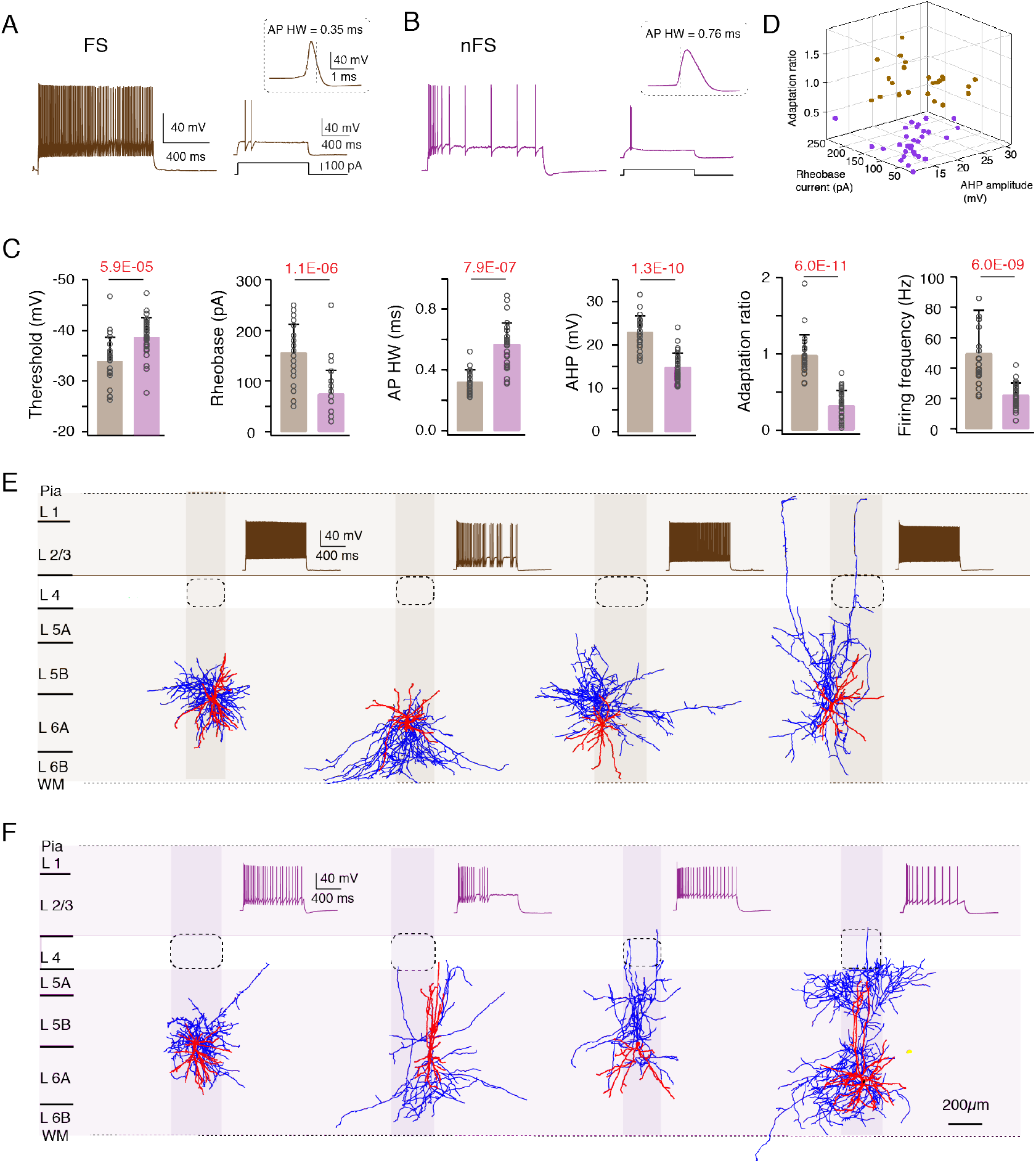
Two major electrophysiological interneuron subgroups in L6A of rat barrel cortex. **(A, B)** Left, representative firing patterns of a L6A FS (A) and a nFS (B) interneuron. Right, responses of a FS and a nFS interneuron to rheobase current injection. Inset showing the first AP in higher magnification. **(C)** Summary data of several electrophysiological properties of L6A interneurons. P values are calculated using Wilcoxon Mann-Whitney U test. **(D)** 3D scatter plot showing the clear separation of FS (n = 23) and nFS (n = 30) interneurons using electrophysiological properties. FS interneurons in brown and nFS interneurons in purple. **(E, F)** Representative morphological reconstructions and the corresponding firing patterns of 4 FS and 4 nFS (F) interneurons. Both FS and nFS interneurons show diverse axonal projection patterns, suggesting that both groups comprise several different interneuron types. Barrels and home columns are indicated in light gray.

GABAergic interneurons in layer 6 of the neocortex show a high degree of morphological diversity (Kumar and Ohana 2008; Bortone et al. 2014; Arzt et al. 2018; Gouwens et al. 2019; Ding et al. 2020). In agreement with previous findings, heterogeneous axonal innervation profiles were identified for L6A FS and nFS interneurons indicating that the firing pattern of L6A interneurons is not tightly correlated with axonal projection patterns (**Fig. 3 E, F**). While some were local interneurons with axonal collaterals confined to deep layers others are inter-laminar projecting interneurons with axons extending to more superficial layers. The axonal projections of most L6A interneuron types were not confined to the borders of the ‘home’ cortical column which in layer 6 are delimited by the so-called L6 ‘infra-barrels’ (Crandall et al. 2017), but innervate also neighbouring ‘barrel columns’ (**Fig. 3 E, F**).

Paired-recordings were performed between presynaptic PCs and postsynaptic interneurons in L6A. 369 potential connections were probed between presynaptic excitatory cells and postsynaptic interneurons. 39 excitatory synaptic connections were detected resulting in a connectivity ratio of 10.6 %. For 30 E→I cell pairs both presynaptic PCs and postsynaptic interneurons were morphologically reconstructed and electrophysiologically analysed, allowing post-hoc identification of connection subtypes in accordance with pre- and postsynaptic cell classes (**Fig. S7**). As for E→E connections with a presynaptic CT PC, CT→interneuron connections were found to be weak (1st EPSP amplitude ranging from 0.02 to 0.36 mV) and unreliable and displayed short-term facilitation (**Fig. 4 A, B**). The functional properties of synaptic connections between CT PCs and FS or nFS interneurons, respectively were not significantly different. However, CT→FS connections tended to show slightly larger mean uEPSP amplitudes and weaker synaptic facilitation (**Fig. 4 C, Tab. 2**). On the other hand, L6A FS and nFS interneurons showed distinct responses to presynaptic stimulation of CC PCs (Fig. 5 A-B). Synaptic connections between CC PC and (postsynaptic) L6A FS interneurons displayed short-term depression and a large mean uEPSP amplitude (1.13 ± 0.78 mV), low CV (0.62 ± 0.26), and low failure rate (20.7 ± 20.6 %). In contrast, CC→nFS connections showed on average a small uEPSP amplitude (0.16 ± 0.18 mV), high CV (1.22 ± 0.28) and high failure rate (59.0 ± 24.3 %) and exhibited short-term facilitation (**Fig. 4 C, Tab. 2**). Similarly, a depression or facilitation of the postsynaptic response can be observed in CCla→FS and CCla→nFS connection, respectively (**Fig. S8**). It is worth noting, however, that both the FS and nFS interneuron show a large mean uEPSP amplitude and a low failure rate in response to presynaptic APs of CCla PC (**Fig. S8**). Statistical comparison of other synaptic properties between different E→I connection types are given in **Tab.2**. Our results suggest that both pre- and postsynaptic cell types govern synaptic characteristics in L6A E→I connections.

**Fig. 4.**
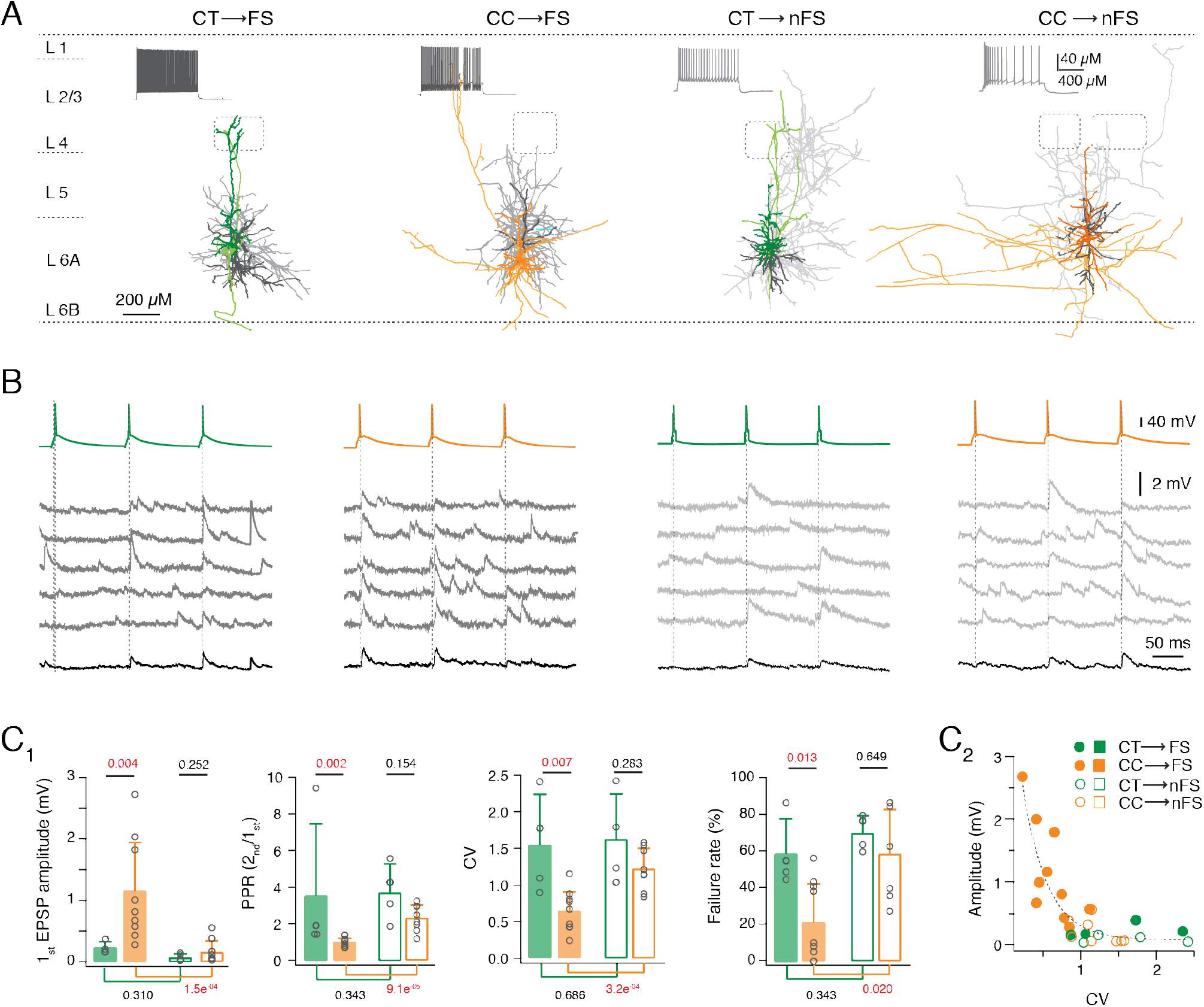
Both pre- and postsynaptic L6A neuron type govern synaptic characteristics of L6A E→I connections. **(A)** Representative morphological reconstructions of a L6A CT→FS, a CC→FS, a CT→nFS and a CC→nFS synaptic connection. Neurons are shown in their approximate laminar location with respect to averaged cortical layers. Presynaptic somatodendritic domain is in a darker, presynaptic axons in a lighter shade, postsynaptic soma and dendrites are in dark gray and postsynaptic axons in light gray. **(B)** Unitary synaptic connections obtained from a CT→FS, a CC→FS, a CT→nFS and a CC→nFS pair. Five consecutive EPSPs (middle) and average EPSP (bottom) are elicited by three consecutive presynaptic APs (top, inter-stimulus interval 100 ms). **(C1)** Summary data of several synaptic properties of L6A E→I connections. P values are calculated using Wilcoxon Mann-Whitney *U* test. **(C2)** First uEPSP amplitude versus CV of L6A E→I connections. Note that the CC→FS connections have large EPSP amplitude and a small CV while the other 3 E→I types are characterised by small mean uEPSP amplitudes and a large CV. Best linear and exponential fits are shown in gray dashed line.

**Fig. 5.**
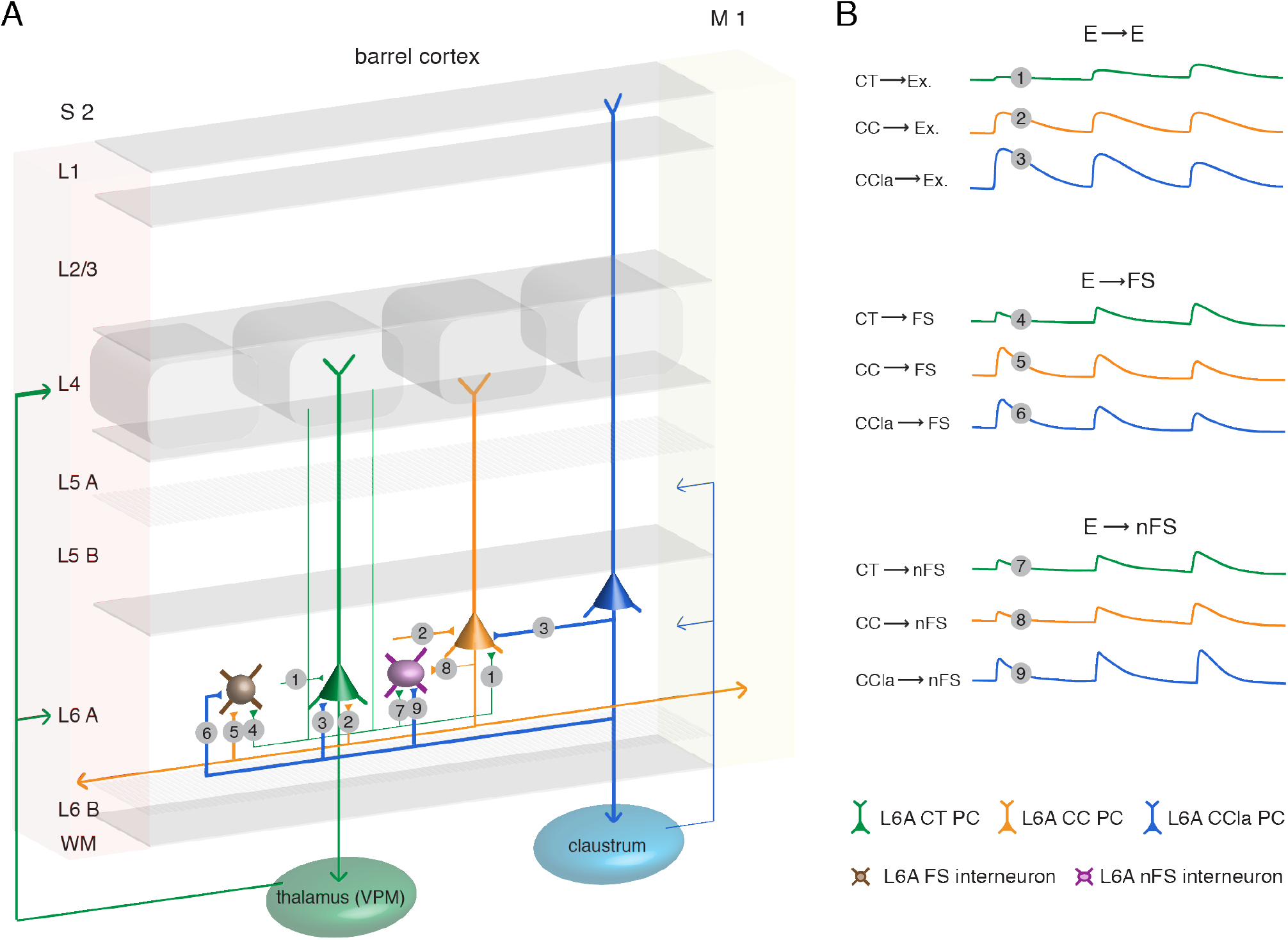
Schematic summary of the excitatory synaptic connections in L6A of rat barrel cortex. **(A)** Synaptic wiring scheme between L6A PCs (E→E) and between L6A PC and interneuron (E→I). The obtained E→E morphological connection types in this study are: CT→CT (no.1), CT→CC (no.1), CC→CT (no.2), CC→CC (no.2), CCla→CT (no.3) and CCla→CC (no.3) connections. The obtained E→I morphological connection types in this study are: CT→FS (no. 4), CC→FS (no.5), CCla→FS (no.6), CT→nFS (no.7), CC→nFS (no.8) and CCla→nFS (no.9) connections. The thickness of axonal projections indicating the strength of synaptic release. WM, white matter; S2, secondary somatosensory cortex; M1, primary motor cortex. **(B)** The 1st uEPSP amplitude and the short term plasticity differ at the different L6A excitatory connection types. CT PCs form weak, facilitating connections with other L6A PCs and interneurons. Excitatory L6A CC PCs connections show no obvious short-term depression or facilitation. CC→FS connections display a large 1st EPSP amplitude with short-term depression but establish weak and facilitating synapses with L6A nFS interneuron. CCla→interneuron connections are strong but CCla→FS connections display short-term depression while CCla→nFS connection show short-term facilitation.

**Tab. 2.**
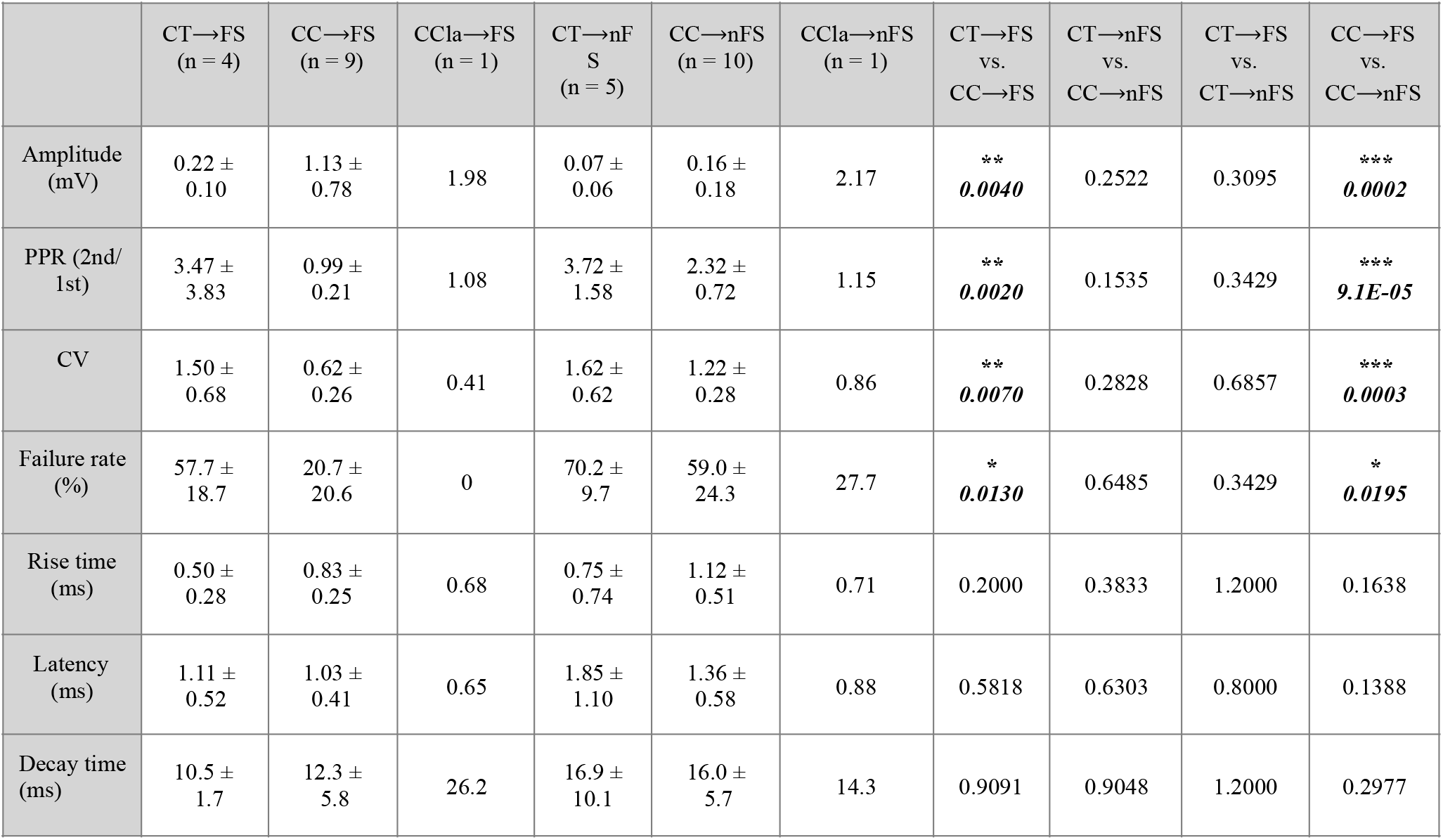
Functional properties of L6A E→I synaptic connections. Italic bold font indicates significant differences; *P < 0.05, **P < 0.01, ***P < 0.001 for Wilcoxon Mann-Whitney *U* test.

|  | CT→FS<br>(n = 4) | CC→FS<br>(n = 9) | CCla→FS<br>(n = 1) | CT→nF<br>S<br>(n = 5) | CC→nFS<br>(n = 10) | CCla→nFS<br>(n = 1) | CT→FS<br>vs.<br>CC→FS | CT→nFS<br>vs.<br>CC→nFS | CT→FS<br>vs.<br>CT→nFS | CC→FS<br>vs.<br>CC→nFS |
| --- | --- | --- | --- | --- | --- | --- | --- | --- | --- | --- |
| Amplitude<br>(mV) | 0.22 ±<br>0.10 | 1.13 ±<br>0.78 | 1.98 | 0.07 ±<br>0.06 | 0.16 ±<br>0.18 | 2.17 | **<br>**0.0040** | 0.2522 | 0.3095 | ***<br>**0.0002** |
| PPR (2nd/<br>1st) | 3.47 ±<br>3.83 | 0.99 ±<br>0.21 | 1.08 | 3.72 ±<br>1.58 | 2.32 ±<br>0.72 | 1.15 | **<br>**0.0020** | 0.1535 | 0.3429 | ***<br>**9.1E-05** |
| CV | 1.50 ±<br>0.68 | 0.62 ±<br>0.26 | 0.41 | 1.62 ±<br>0.62 | 1.22 ±<br>0.28 | 0.86 | **<br>**0.0070** | 0.2828 | 0.6857 | ***<br>**0.0003** |
| Failure rate<br>(%) | 57.7 ±<br>18.7 | 20.7 ±<br>20.6 | 0 | 70.2 ±<br>9.7 | 59.0 ±<br>24.3 | 27.7 | *<br>**0.0130** | 0.6485 | 0.3429 | *<br>**0.0195** |
| Rise time<br>(ms) | 0.50 ±<br>0.28 | 0.83 ±<br>0.25 | 0.68 | 0.75 ±<br>0.74 | 1.12 ±<br>0.51 | 0.71 | 0.2000 | 0.3833 | 1.2000 | 0.1638 |
| Latency<br>(ms) | 1.11 ±<br>0.52 | 1.03 ±<br>0.41 | 0.65 | 1.85 ±<br>1.10 | 1.36 ±<br>0.58 | 0.88 | 0.5818 | 0.6303 | 0.8000 | 0.1388 |
| Decay time<br>(ms) | 10.5 ±<br>1.7 | 12.3 ±<br>5.8 | 26.2 | 16.9 ±<br>10.1 | 16.0 ±<br>5.7 | 14.3 | 0.9091 | 0.9048 | 1.2000 | 0.2977 |

## Discussion

It has been suggested that in the neocortex, the laminar position of a neuronal cell body accounts for the differences in connection probability and short-term synaptic dynamics (Lefort and Petersen 2017; Seeman et al. 2018; Frandolig et al. 2019). However, for an in-depth understanding of the organization of intra-laminar connectivity in the neocortex a thorough classification of the neuronal cell types in a given cortical layer is crucial (Kiritani et al. 2012; Kawaguchi 2017; Anastasiades et al. 2019; Whitesell et al. 2020). In previous studies of L6A microcircuits CCla PCs were often overlooked (West et al. 2006; Crandall et al. 2017; Sundberg et al. 2018; Frandolig et al. 2019), probably because of their low density and their differential laminar distribution in various cortical areas (Gutierrez-Lbarluzea et al. 1999; Wang et al. 2017). Here, we identified three distinct types of L6A PCs based on both their electrophysiological and anatomical features; these L6A PC types were named CT, CC and putative CCla PCs based on their putative axonal targets. Furthermore, we were able to show that these L6A PC subpopulations establish excitatory synaptic connections with very distinct dynamic properties.

### Cortico-thalamic PCs

In deep layers of the neocortex, principal neurons with projections confined to the telencephalon preferentially generate depressing EPSPs whereas ET-projecting pyramidal cells tend to display short-term facilitation (West et al. 2006; Le Be et al. 2007; Morishima et al. 2011; Cotel et al. 2018). In accordance with this view, our results showed that presynaptic CT PCs projecting to the ventral posterior medial nucleus (VPM) form excitatory connections that display strong short-term facilitation (Killackey and Sherman 2003), while those formed by presynaptic intracortical CC PCs and putative CCla PCs display only weak facilitation or depression. Apart from a population of remnant subplate neurons in layer 6 (Marx et al. 2017; Hoerder-Suabedissen et al. 2018), L6 CT PCs may be the earliest neuron class to populate the developing neocortex (Auladell et al. 2000). There is evidence that CC PCs in in layer 6 of secondary somatosensory cortex are born later than CT PCs (Arimatsu et al. 1999; Arimatsu and Ishida 2002). With developmental maturation, glutamatergic synapses turn to short-term facilitation concomitant with a reduction in synaptic release probability (Oswald and Reyes 2008; Feldmeyer and Radnikow 2009). Thus, it is conceivable that the short-term facilitation of E→E connections reflects the degree of maturation of a synapse type; synapses formed by presynaptic CC and CCla PCs are in a more immature state than L6A CT PCs and thus display more short-term depression (**Fig. 5**). Moreover, synaptotagmin-7 and synapsin I have been shown to have important functions in short-term synaptic facilitation at CT synapses (Nikolaev and Heggelund 2015; Jackman et al. 2016). If these molecules are also present at presynaptic terminals of *intracortical* connections established by CT PCs this may explain - at least in part - EPSP facilitation at these synapses.

In accordance with previous studies (West et al. 2006; Cotel et al. 2018), we found that short-term facilitation is an important feature for the identification of L6A connections with a presynaptic CT PC regardless of the postsynaptic neuron type (**Fig. 5**). There are two known subgroups of L6 CT PCs in rat somatosensory cortex. Some lower L6A CT PCs project to both the ventral posterior medial nucleus (VPM) and the posterior medial nucleus (PoM) of thalamus while the other CT PCs are biased on the upper L6A and projects only to VPM (Zhang and Deschenes 1997; Killackey and Sherman 2003; Chevee et al. 2018). A recent study showed that unlike CT PCs projecting to VPM alone, those projecting to both VPM and PoM establish strong and depressing synapses with L6A PV positive FS interneurons (Frandolig et al. 2019). Here, we did not find this specific connection type, probably because L6A CT PCs projecting to both VPM and PoM have a lower connection probability than the VPM-projecting L6A CT PC subtype (Frandolig et al. 2019). In addition, optogenetic stimulation of CT PCs resulted in facilitating synaptic responses in excitatory cells, FS and nFS interneurons of layers 4 and 5 (Kim et al. 2014), suggesting that intra- and inter-laminar connections with a presynaptic CT PC share common features.

### Cortico-cortical PCs

Compared to L6A CT and CCla PCs, CC PCs showed a higher connection probability with both other L6A PCs and interneurons. A fraction of the L6A CC PCs (~10%) was found to form reciprocal synaptic connections with one another; however, for CT or CCla PCs reciprocal connections were not detected. This suggests that intra-laminar feed-back excitation in layer 6A may be a neuronal cell-type-specific property (Morishima et al. 2011). Positive feedback excitation can drive a prolonged response to brief stimuli, thus maintaining burst activity (Grillner and Graybiel 2006; Li et al. 2006). During sensory processing, feedback excitation increases the sensitivity of CC PCs to thalamic inputs, so that sensory signals can spread quickly and widely spread to the neighbouring barrel columns and even to other cortical areas (Douglas et al. 1995; Lim et al. 2012). On the other hand, studies using micro-iontophoretic injections demonstrated that the long, horizontally projecting axons of CC PCs form reciprocal connections between the somatosensory barrel cortex, the secondary somatosensory, the primary motor, and the perirhinal cortices (Izraeli and Porter 1995; Zhang and Deschenes 1998; for a review see: Aronoff et al. 2010). Here, we also detected several inter-column connections formed by CC PCs with a lateral somatic distance of more than 200 μm. This parallel organization of corticocortical connections in deep layers allows a fast convergence of thalamocortical inputs and in turn drives the reliable sensory responses (Egger et al. 2020).

Apart from the horizontally projecting axons in layer 6, axon collaterals of L6 CC PCs also project extensively to superficial layers where they are likely to contact the apical tufts of thick-tufted L5B PCs (Egger et al. 2020). When proximal synaptic inputs to the basal dendrites of L5B PCs induce a somatic back-propagating APs, coincident synaptic input from L6A CC PCs to the apical tuft may sum with the back-propagating AP to trigger a dendritic calcium spike, a mechanism that is involved in the association of sensory inputs, perception, and learning (Larkum et al. 1999; Larkum and Zhu 2002; Larkum 2013; Takahashi et al. 2016). Thus, L6A CC PCs may have important influence on synaptic integration of L5 PCs, amplifying the response to the thalamocortical inputs while maintaining the neuronal selectivity (Hay and Segev 2015).

### Putative cortico-claustral PCs

In rodents, almost all cortical areas have been found to provide synaptic input to the claustrum; in turn, the claustrum has axonal projections back to all ipsilateral cortical areas and to several contralateral cortical areas (Zakiewicz et al. 2014; Atlan et al. 2017). Although the claustrum is widely connected with cortex, the density of cortico-claustral inputs varies considerably between different species and also different cortical areas (Zingg et al. 2018; Smith et al. 2019a; Jackson et al. 2020), so that functional role of the claustrum is not very well understood. It has been shown that the claustrum responds to stimuli of different sensory modalities and is thereby involved in processing sensory information (Remedios et al. 2010, 2014; Atlan et al. 2018). In barrel cortex cortico-claustral and claustro-cortical axonal projections have been identified by retrograde tracing; they were found to originate or terminate, respectively in deep layers (Zhang and Deschenes 1998; Atlan et al. 2017). CCla PCs described here in layer 6A of rat barrel cortex are a homogeneous subpopulation both with respect to morphology and electrophysiology. They have ascending apical dendrites ending in layer 1 and widely expanding basal dendrites within layer 6, morphological features that are highly similar to those of CCla PCs in cat and rat primary visual cortex (Katz 1987; Cotel et al. 2018). In layer 6 of rat prefrontal cortex, a high percentage of PCs (approximately 40%) show the tall, wide dendritic morphology suggesting that this morphological subtype exist in many different cortical regions (Van Aerde and Feldmeyer 2015).

Another potential target of IT L6A PCs is the striatum as part of the basal ganglia. The striatum receives glutamatergic inputs directly from various cortical areas as well as the (extratelencephalic) thalamus making it well suited for integrating sensory information. Striatal neurons respond to sensory input from different modalities including tactile, auditory, and visual input (Reig and Silberberg 2014). Pyramidal cells projecting to the striatum are distributed throughout layer 2 to 6 (Anastasiades et al. 2019; Zhang et al. 2019). Because the density of corticostriatal PCs is particularly high in cortical layer 5 most studies have focussed on these neurons (Shepherd 2013). Nevertheless, a correlated description of the morphological, electrophysiological, and synaptic characteristics of L6 corticostriatal PCs is so far not available.

It is of note that the putative L6A CCla PCs identified here show a higher membrane excitability and stronger synaptic release than other L6A PC populations. This suggests that although they form only a small fraction in layer 6A, CCla PCs are actively involved in local circuits. L6A CCla PCs preferentially innervate CC rather than CT PCs forming strong and stable connections. This suggests that they may contribute in coordinating a wide-ranging network between different cortical regions. Furthermore, neocortical nFS interneurons appear to establish weak synaptic connections with neighbouring PCs that show short-term facilitation, resulting in a delayed recruitment of inhibition via these interneurons (Helmstaedter et al. 2008; Caputi et al. 2009). This was also observed with CT→nFS and CC→nFS connections (**Fig. 5**). However, the CCla→nFS connection we recorded was also found to be strong and reliable, suggesting that the synaptic microcircuitry formed by L6A CCla PCs is uniquely dominant despite the low density of these neurons.

In conclusion, we have demonstrated that the excitatory synaptic microcircuitry in layer 6A of rat barrel cortex is highly specific for the excitatory neuronal cell type, with important implications for understanding intracortical network and subcortical output of layer 6 as well as there feedback and feed-forward projections. Our study provides novel data necessary to obtain a more complete and coherent picture of L6 microcircuitry.

## Acknowledgement

We would like to thank Werner Hucko for excellent technical assistance and Dr. Karlijn van Aerde for custom-written macros in Igor Pro software. We are also grateful to Dr. Manuel Marx for sharing preliminary data for this project. We thank Carmen Kapitel and Jawad Jawadi for help in Neurolucida reconstructions. We warmly thank Dr. Chao Ding for helpful discussion.

