## supplementary materials for "Layer 6A pyramidal cells subtypes form synaptic microcircuits with distinct functional and structural properties"

Containing 1 table and 8 figures with legends

Danqing Yang^1,^*, Guanxiao Qi^1,^* and Dirk Feldmeyer^1,2,3,★^

^1^ Research Centre Juelich, Institute of Neuroscience and Medicine (INM-10), Leo-Brandt-Strasse, Juelich, Germany

^2^ RWTH Aachen University Hospital, Pauwelsstrasse 30, Aachen, Germany

^3^ Jülich Aachen Research Alliance, Translational Brain Medicine (JARA Brain), Aachen, Germany

* These authors have contributed equally to this work

^★^Corresponding author

Dirk Feldmeyer

Research Centre Juelich

Institute of Neuroscience and Medicine 10

Section ‘Function of Neuronal Microcircuits’

Leo-Brandt-Strasse

52425 Juelich

Germany

**Tab. S1 Morphological and electrophysiological properties of L6A pyramidal cells.**

Italic bold font indicates significant differences; *P < 0.05, **P < 0.01, ***P < 0.001 for Wilcoxon-Mann-Whitney *U* test.

|  | CT | CC | CCla | CT vs. CC | CT vs. CCla | CC vs. CCla |
| --- | --- | --- | --- | --- | --- | --- |
| Morphological properties (CT n = 11; CC n = 16; CCla n = 14) | | | | | | |
| Nr. of basal dendrites | 6.6 ± 1.6 | 6.3 ± 1.8 | 3.6 ± 0.9 | 0.6789 | ******4.5E-06*** | ******6.0E-06*** |
| total length of basal dendrites (µm) | 1899 ± 447 | 3770 ± 1277 | 3115 ± 950 | ******7.7E-05*** | *****0.0011*** | 0.1934 |
| average length of basal dendrites (µm) | 295 ± 59 | 607 ± 170 | 917 ± 358 | ******1.8E-06*** | ******4.5E-07*** | *****0.0051*** |
| dendritic horizontal feldspan (µm) | 231 ± 58 | 386 ± 57 | 733 ± 146 | ******1.0E-05*** | ******4.5E-07*** | ******6.8E-09*** |
| dendritic vertical fieldspan (µm) | 773 ± 184 | 731 ± 163 | 1546 ± 237 | 0.3356 | ******4.5E-07*** | ******1.4E-08*** |
| axonal length  (µm) | 5502 ± 2189 | 16356 ± 4081 | 10224 ± 4648 | ******3.1E-07*** | *****0.0090*** | *****0.0021*** |
| axonal horizontal fieldspan (µm) | 529 ± 240 | 1545 ± 541 | 1101 ± 479 | ******6.1E-07*** | ******0.0003*** | *****0.0069*** |
| relative soma depth (%) | 78.5 ± 4.1 | 66.6 ± 6.5 | 69.4 ± 5.2 | ******0.0004*** | ******0.0001*** | 0.1600 |
| Electrophysiological properties (CT n =15; CC n = 23; CCla n = 22) | | | | | | |
| resting membrane potential (mV) | -69.2 ± 4.4 | -71.1 ± 5.7 | -69.6± 3.4 | 0.2266 | 0.7952 | 0.2269 |
| input resistance  (MΩ) | 235.1 ± 79.5 | 151.7 ± 35.1 | 281.3 ± 73.2 | ******0.0001*** | 0.0804 | ******7.1E-10*** |
| rheobase current (pA) | 102.0 ± 42.6 | 132.7 ± 39.4 | 62.5 ± 23.4 | ****0.0180*** | *****0.0014*** | ******6.2E-08*** |
| AP latency  (ms) | 129.7 ± 32.3 | 177.6 ± 54.2 | 203.3 ± 81.0 | ******0.0001*** | ******7.3E-05*** | 0.2942 |
| AP threshold  (mV) | -35.1 ± 3.5 | -35.2 ± 3.5 | -39.8 ± 3.3 | 0.8828 | ******0.0007*** | ******6.0E-05*** |
| ISI_2_/ISI_10_ | 0.76 ± 0.16 | 0.43 ± 0.23 | 0.46 ± 0.20 | ******2.3E-05*** | ******0.0002*** | 0.7937 |
| 1_st_ AP amplitude (mV) | 93.1 ± 8.0 | 91.9± 7.0 | 102.2 ± 7.2 | 0.6580 | ******0.0007*** | ******2.6E-05*** |
| 2_nd_ AP amplitude (mV) | 80.7 ± 11.6 | 61.5 ± 11.3 | 92.1 ± 10.3 | ******1.5E-05*** | *****0.0046*** | ******5.5E-11*** |
| 1_st_ AP half-width (ms) | 0.78 ± 0.10 | 0.91 ± 0.14 | 0.72 ± 0.15 | *****0.0043*** | 0.1367 | ******7.3E-05*** |
| 2_nd_ AP half-width (ms) | 0.91 ± 0.10 | 1.44 ± 0.38 | 0.88 ± 0.21 | ******5.4E-08*** | 0.5457 | ******2.0E-07*** |

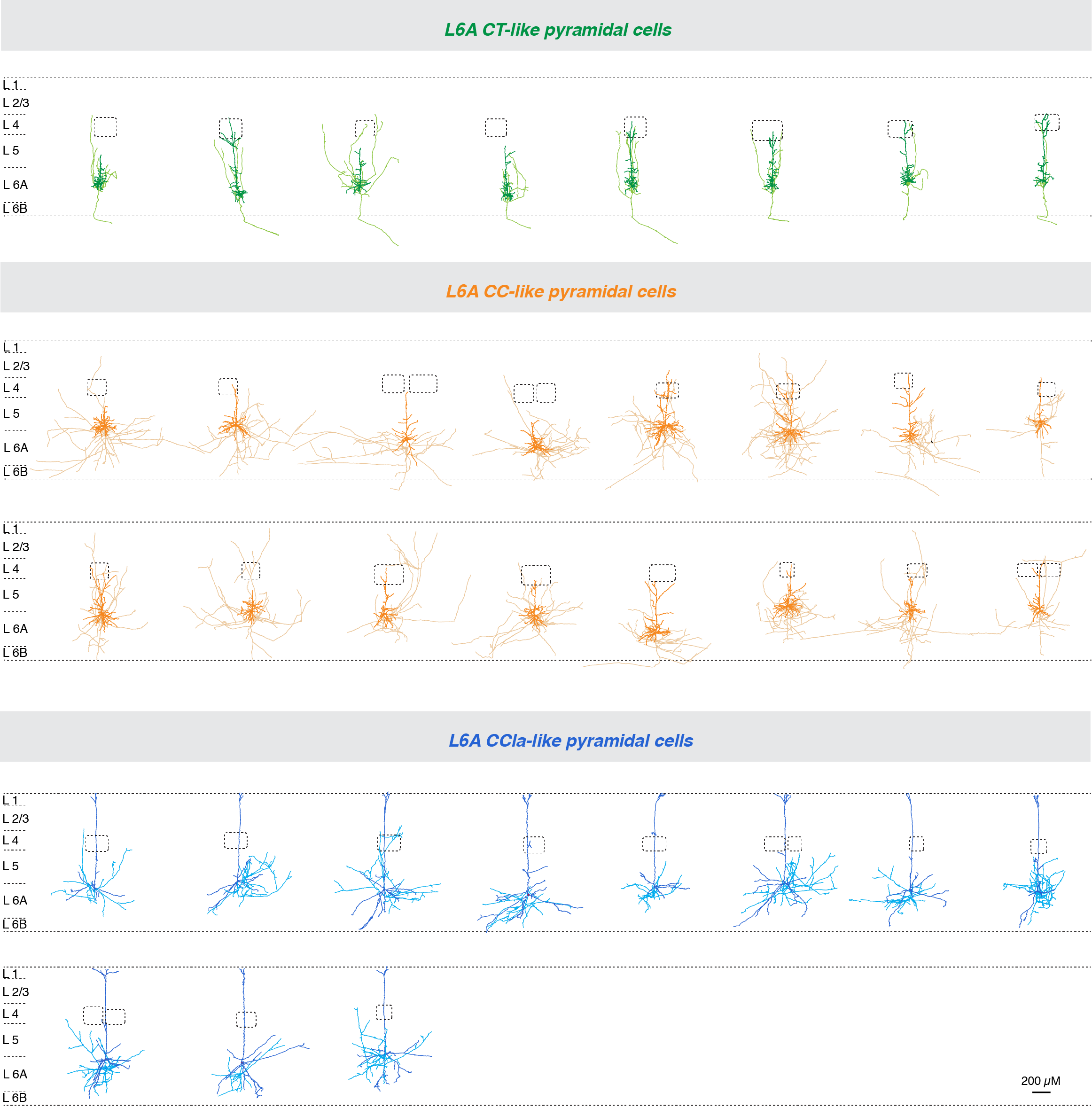

**Fig. S1 Neurolucida reconstructions of L6A PCs.**

Individual morphological reconstruction of the three subtypes of L6A PCs. Neurons are shown in their approximate laminar location with respect to averaged cortical layers. CT-like PCs are in blue, CC-like pyramidal in orange and CCla-Like PCs in blue. The somatodendritic domain is given in a darker, the axons in a lighter shade.

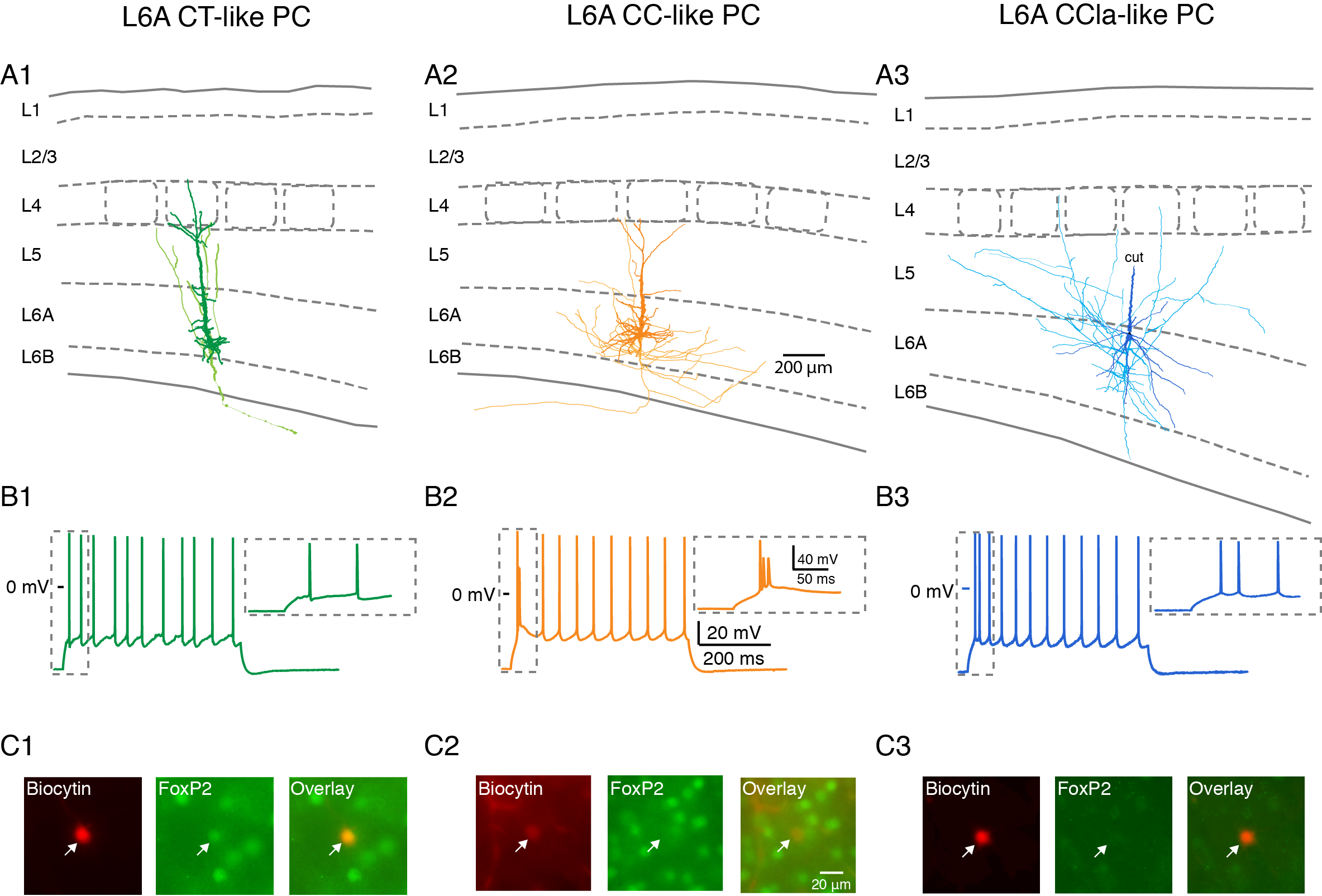

**Fig. S2 Three representative examples of morphologically and electrophysiologically identified L6A PCs which show different FoxP2 expression patterns.**

(A) Morphological reconstructions of a CT (A1), CC (A2) and a putative CCla (A3) L6A PC. Same color code as in Fig. S1. The Somatodendritic domains are shown in dark and axons in light shade. Note that in panel A3 even the apical dendrite has been truncated before it reaches the pial surface, the identity of putative CCla L6A PC could still be deduced by their somatic location and enlarged basal dendrites.

(B) Corresponding firing patterns of the CT (B1), CC (B2) and CCla PC (B3) from (A) are shown.

(C) The neurons shown in (A) were recorded using whole-cell patch-clamp technique and simultaneously filled with biocytin and the fluorescent dye Alexa 594 (red). Antibody labeling was performed to test for the expression of FoxP2 (green). Alexa 594 was used to mark the location of the recorded neuron in the slice.

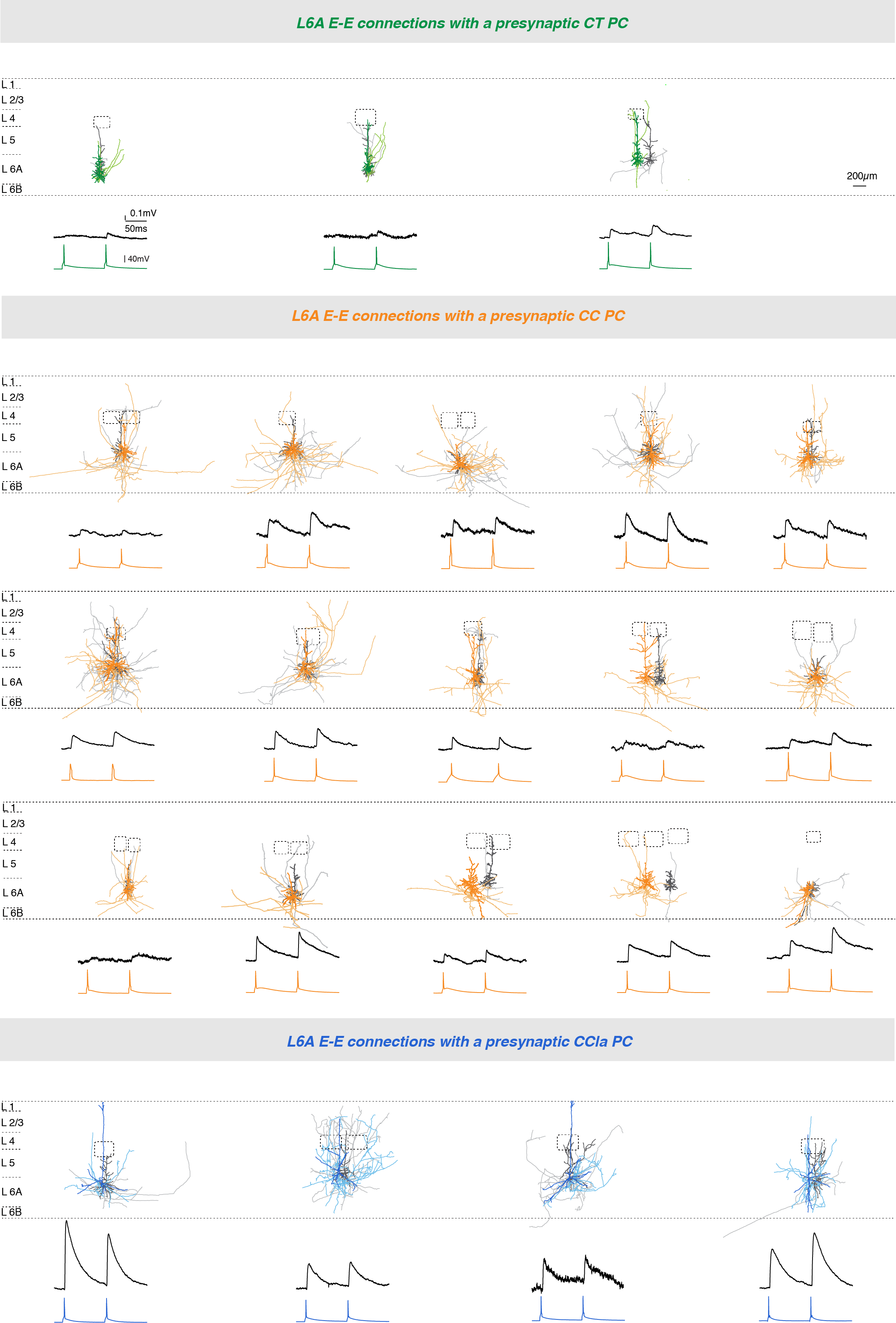

**Fig. S3 Neurolucida reconstructions and electrophysiological recordings of L6A E**⟶**E connections.**

Top, Individual morphological reconstruction of the three subtypes of L6A E⟶E connections. Neurons are shown in their approximate laminar location with respect to averaged cortical layers. Soma and dendrites of the presynaptic PC is given in dark green, orange, or blue, its axons in light green, orange, or blue, respectively; the somatodendritic domain of the postsynaptic PC is dark gray and postsynaptic axons in light gray. Bottom, averaged EPSPs in response to presynaptic APs (10 Hz) obtained from the same E⟶E cell pair shown above.

**Fig. S4 Paired pulse behaviour of L6A E**⟶**E connections.**

(A, B C) Left, representative morphological reconstructions of a L6A CT⟶CT, a CC⟶CC and a CCla⟶CC cell pair. Soma and dendrites of the presynaptic PC are given in a darker while presynaptic axons in a lighter shade, po
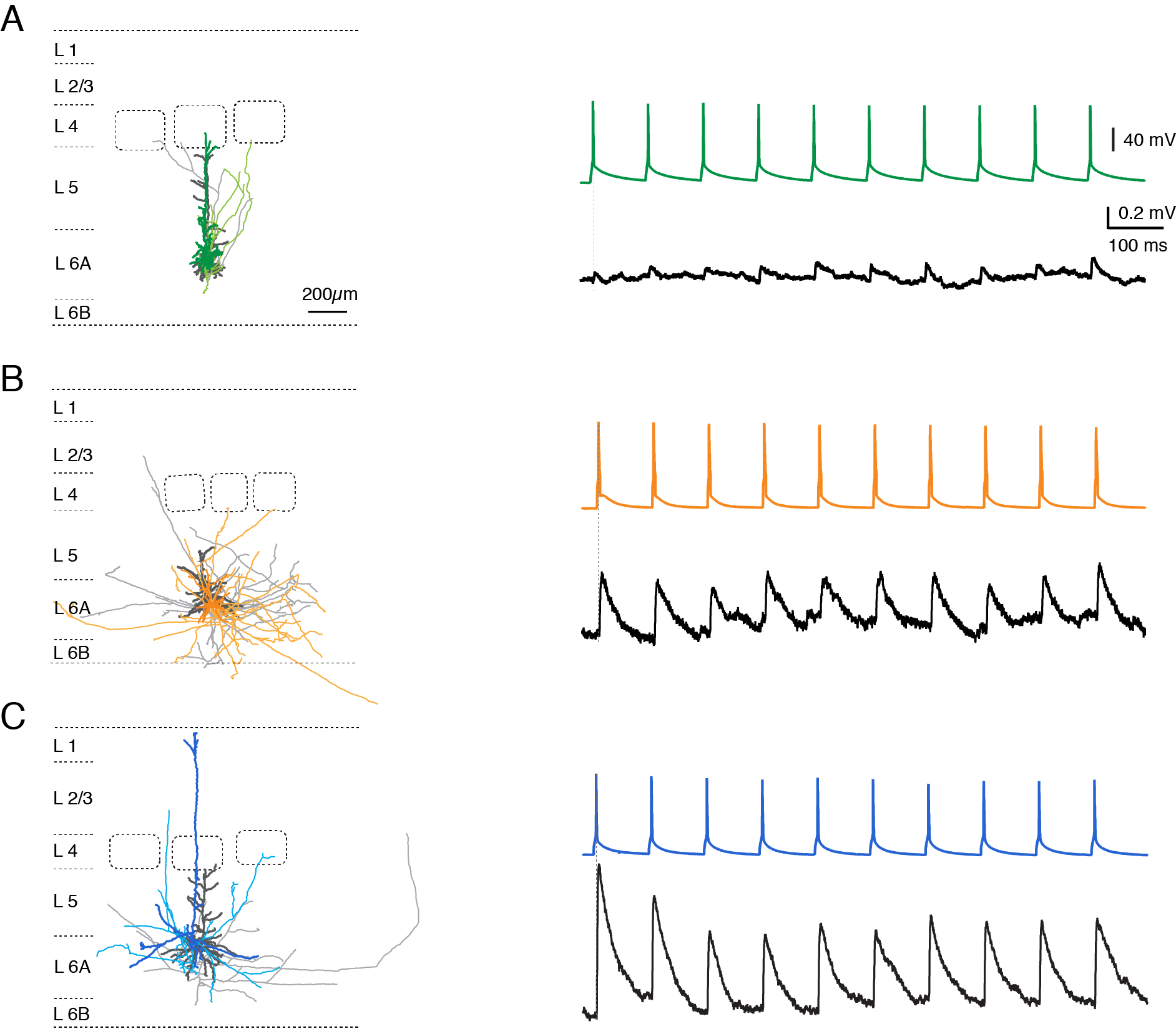
stsynaptic somatodendrites in dark gray and postsynaptic axons in light gray. Color of the different L6 PC subclasses are as in Fig. S1. Right, averaged uEPSPs (bottom) in response to trains of ten consecutive APs (top, 10 Hz) in a presynaptic CT (A), CC (B) and CCla PC (C).

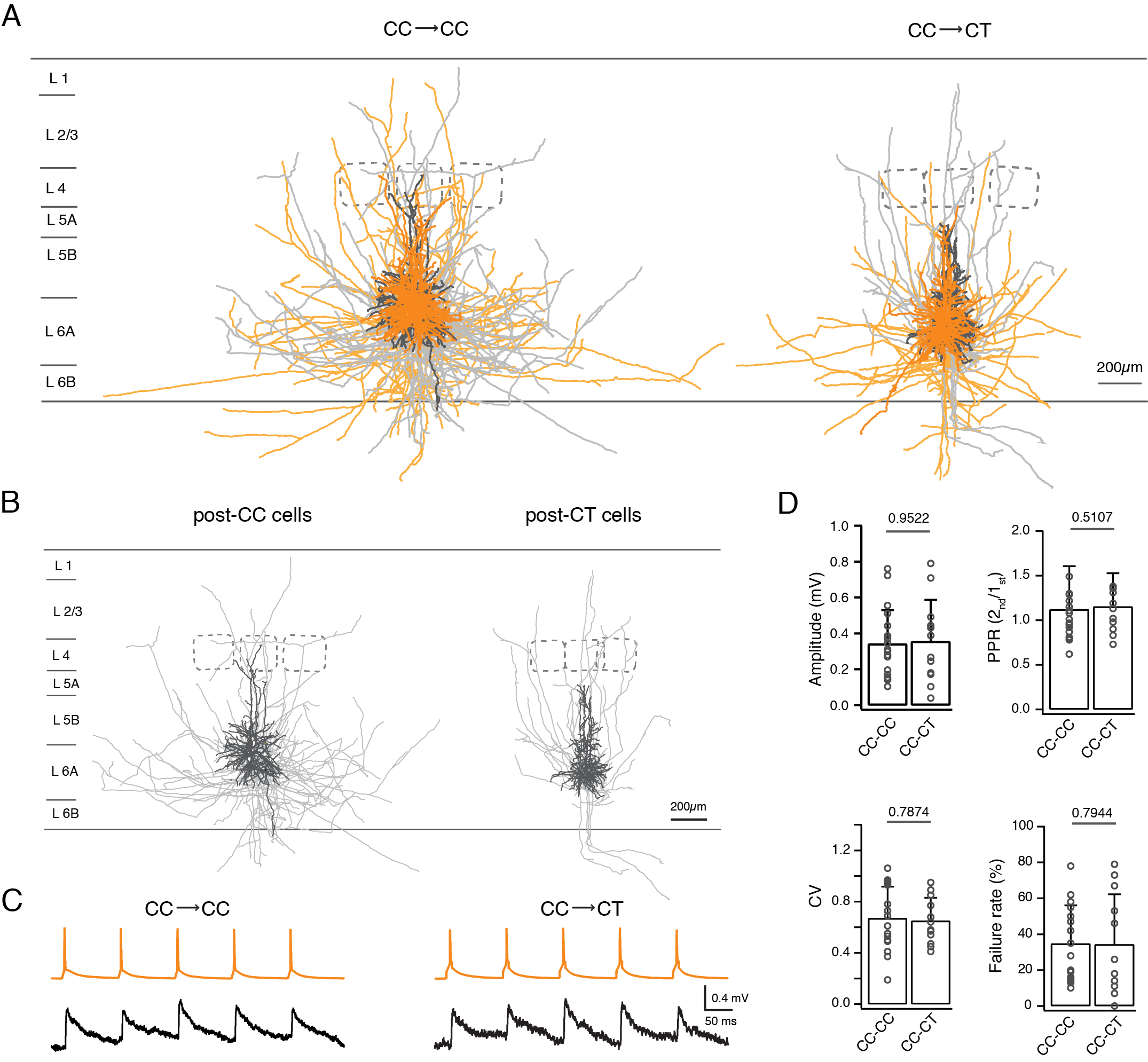

**Fig. S5 Comparison of two types of E**⟶**E connetcions formed by presynaptic CC PCs.**

(A) Overlay of five reconstructions of CC⟶CC (left) and CC⟶CT (right) cell pairs. Presynaptic soma and dendrites are in dark orange, the presynaptic axons in light orange; postsynaptic soma and dendrites are in dark gray and postsynaptic axons in light gray. (B) Somatodendritic and axonal domains of the postsynaptic CC (left) and CT (right) PCs from the same cell pairs as in (A). (C) Averaged postsynaptic EPSPs (bottom) recorded in a CC (left) and CT (right) PC in response to 5 train spikes (top, 10 Hz) of presynaptic CC PCs. (D) Histograms comparing the amplitude of the 1st EPSP, PPR, CV and failure rate for CC⟶CC (n = 25) and CC⟶CT (n = 11) connections. P values are calculated using the Wilcoxon Mann-Whitney U test.

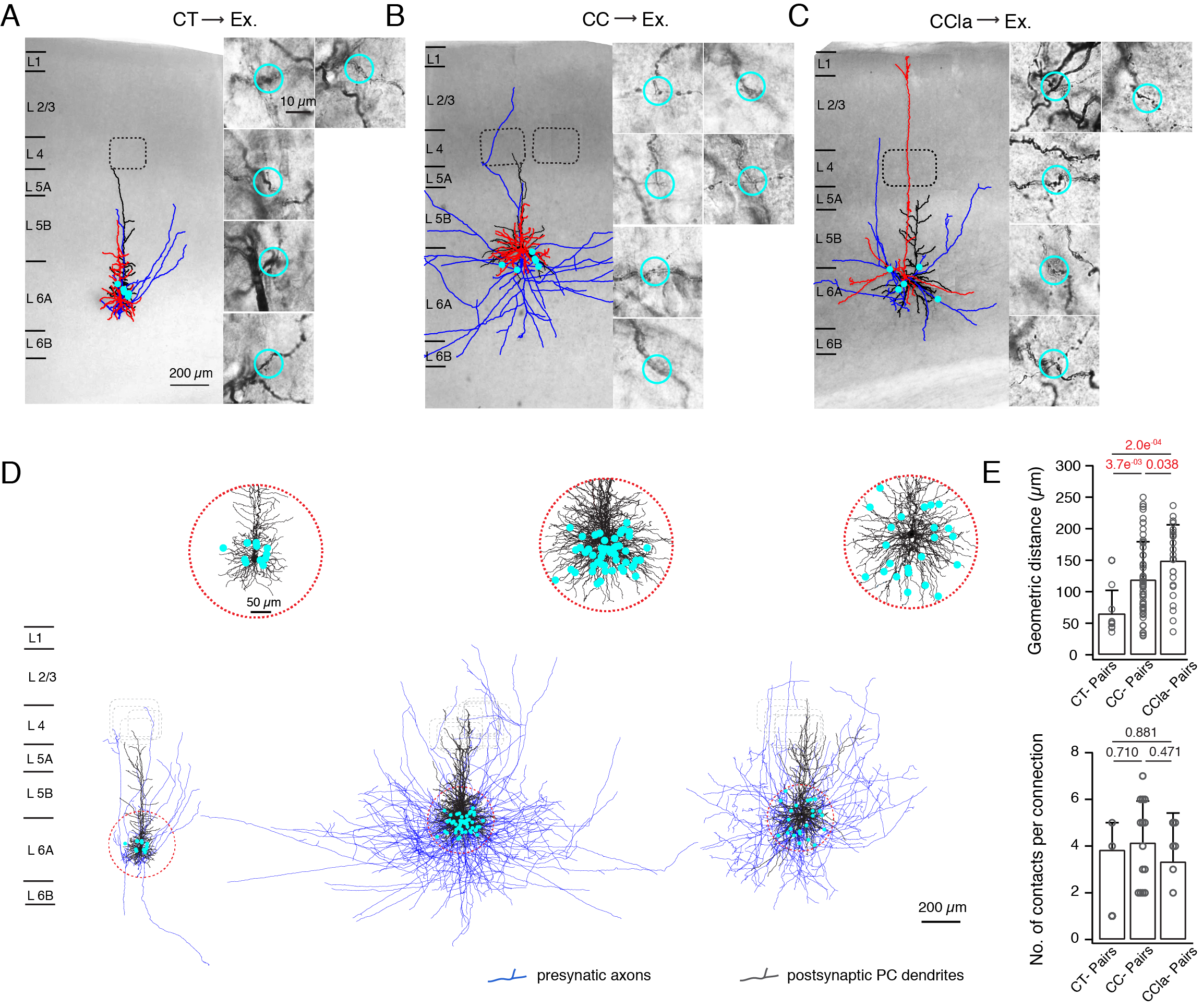

**Fig. S6 L6A The spatial distribution of putative synaptic contacts between E**⟶**E connections depend on the presynaptic L6A PC subtypes.**

(A, B, C) Left, Photomicrograph of exemplary biocytin-labeled CT⟶CT (A), CC⟶CC (B), and CCla⟶CC (C) PC pairs superimposed by the respective morphological reconstructions. Presynaptic soma and dendrites, red; presynaptic axons, blue; postsynaptic soma and dendrites, black. Barrels are indicated by dashed lines. Right, high power photomicrographs of individual light-microscopically identified putative synaptic contacts. Putative contacts are marked by either by light blue dots (left) or circles (right). Synaptic contacts at higher magnification are shown at the right.

(D) Presynaptic axon and postsynaptic soma and dendrites of the three L6A E⟶E connection subtypes. Putative synaptic contacts are marked by red dots. Somatodendritic domain were aligned with respect to the soma position. Insets, higher magnification of the marked regions display the difference in distribution of synaptic contacts among three connection subtypes.

(E) Summary data of morphological properties of L6A E⟶E connections with presynaptic CT (green, n = 4), CC (orange, n = 14) and CCla (blue, n = 6) PCs. P values are calculated using the Wilcoxon Mann-Whitney U test.

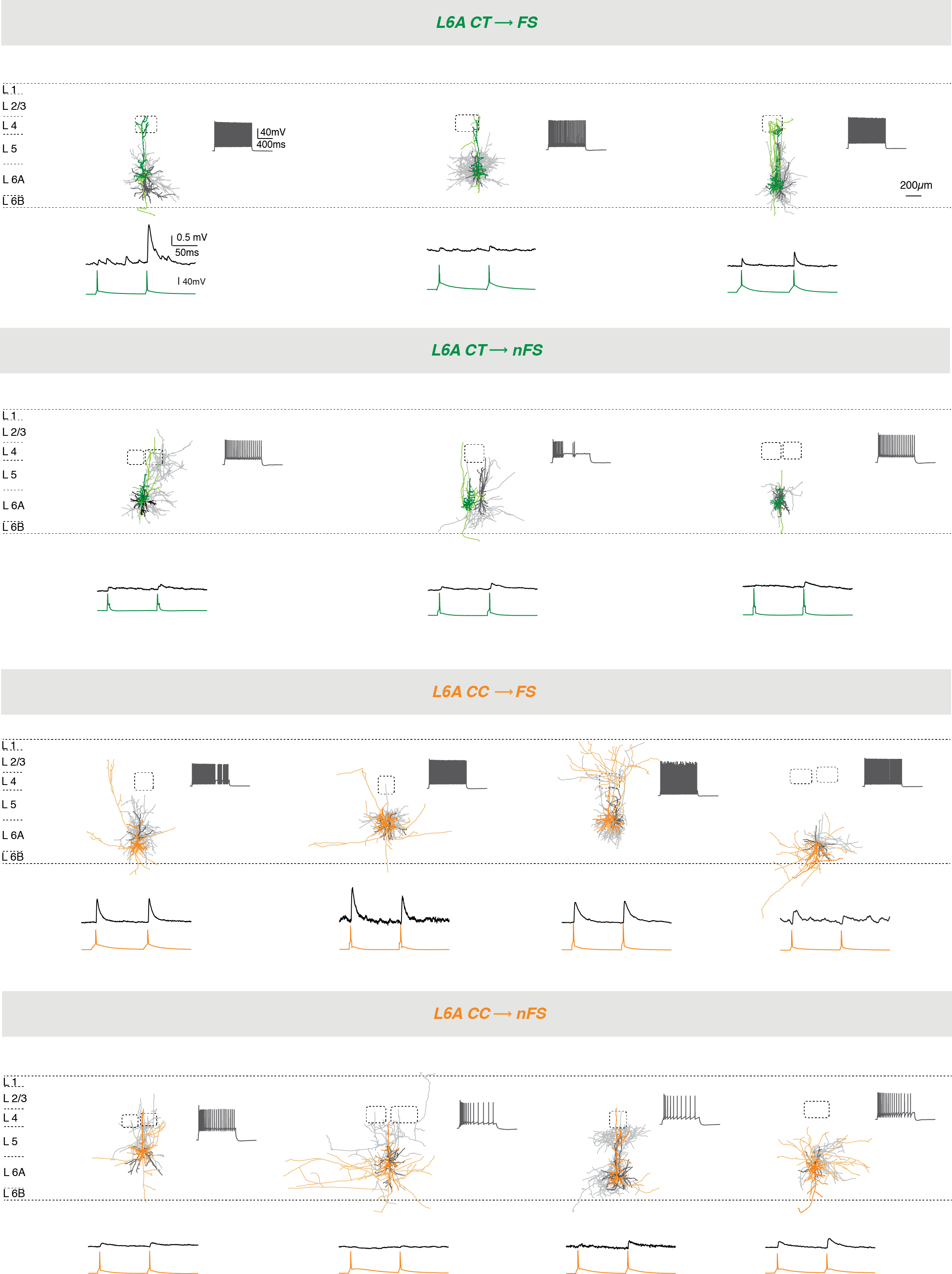

**Fig. S7 Neurolucida reconstructions and electrophysiological recordings of L6A E**⟶**I connections.**

Top, Individual morphological reconstruction of the four subtypes of L6A E⟶I connections. Neurons are shown in their approximate laminar location with respect to averaged cortical layers. Presynaptic somatodendritic domain is in a darker, presynaptic axons in a lighter shade, postsynaptic soma and dendrites are in dark gray and postsynaptic axons in light gray. Inset shows the corresponding firing pattern of postsynaptic interneuron. Bottom, averaged EPSPs in response to presynaptic APs (10 Hz) obtained from the same E⟶I cell pair shown above.

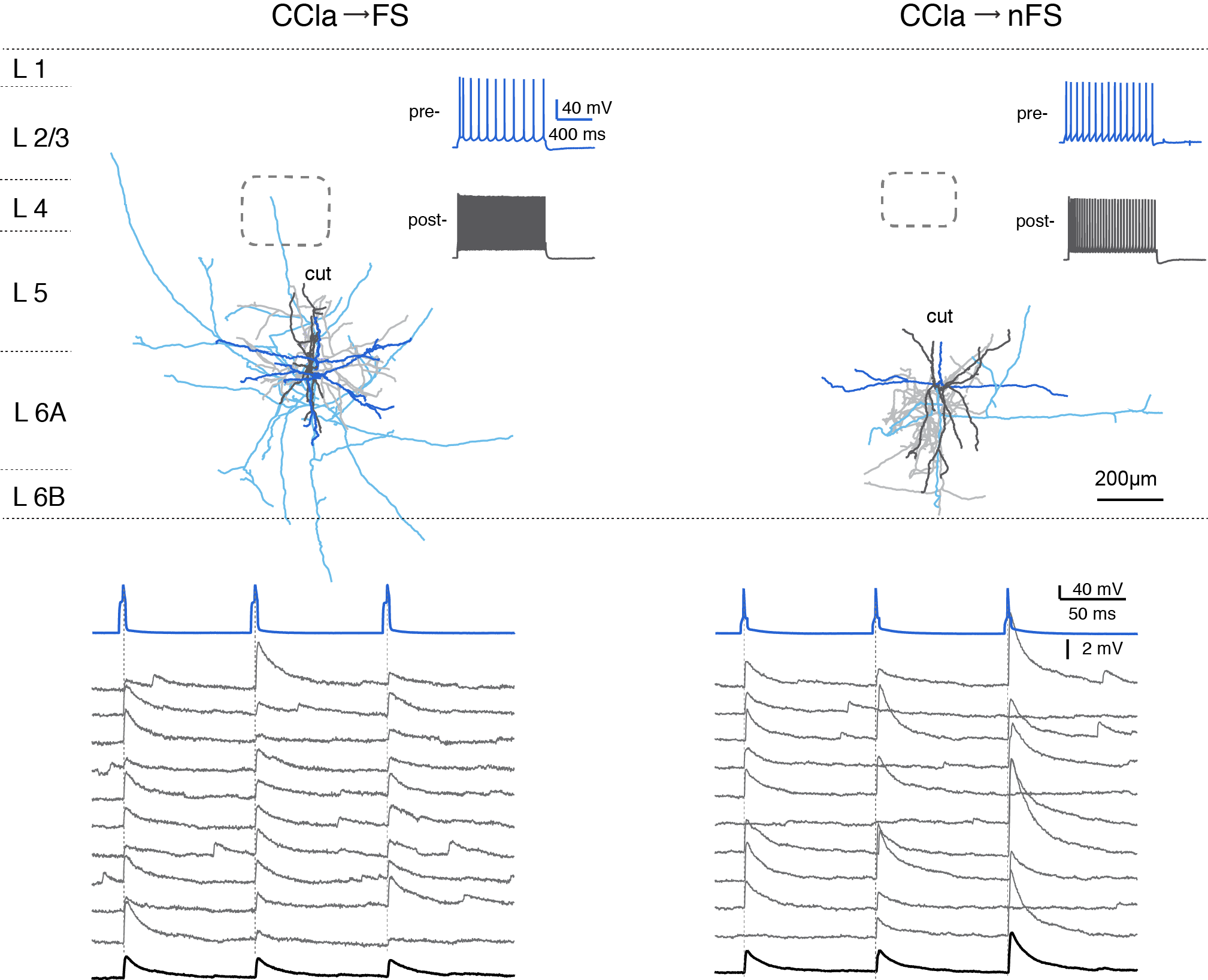

**Fig. S8 Two types E**⟶**I connections formed by presynaptic putative CCla PCs.**

In these pairs CCla PCs were identified based on their location in upper layer 6A, the broad basal dendritic field and their firing pattern (s. also Fig. 1 off the main manuscript). Top, morphological reconstructions of a L6A CCla⟶FS and a CCla⟶nFS cell pair. The soma and dendrites of the presynaptic neurons are given in a dark blue, the presynaptic axons in light blue, postsynaptic soma and dendrites in dark gray and postsynaptic axons in light gray. Insets showing the corresponding firing patterns of pre- and postsynaptic neurons. Bottom, corresponding EPSPs obtained from the shown CCla⟶FS and CCla⟶nFS pair. Ten consecutive EPSPs (middle, gray) and avergae EPSP (bottom, black) are elicited by presynaptic APs (top, blue).
